# Hepatic HKDC1 Deletion Alleviates Western Diet-Induced MASH in Mice

**DOI:** 10.1101/2024.11.26.625530

**Authors:** Kai Xu, Irene Covila-Corona, María Dolores Frutos, María Ángeles Núñez-Sánchez, Dhruvi Makhanasa, Pratham Viral Shah, Grace Guzman, Bruno Ramos-Molina, Medha Priyadarshini, Md. Wasim Khan

**Author notes:** Authors have equal contribution (Kai Xu, Irene Covila-Corona).

## Abstract

The global prevalence of Metabolic dysfunction-associated steatohepatitis (MASH) has been rising sharply, closely mirroring the increasing rates of obesity and metabolic syndrome. MASH exhibits a strong sexual dimorphism where females are affected with more severe forms after menopause. Hexokinase domain-containing protein 1 (HKDC1) has recently been recognized for its role in liver diseases, where its expression is minimal under normal conditions but significantly increases in response to metabolic stressors like obesity and liver injury. This selective upregulation suggests HKDC1’s potential specialization in hepatic glucose and lipid dysregulation, linking it closely to the progression of MASLD and MASH. This study aims to clarify the role of HKDC1 in Western diet-induced MASH in female mice by examining its impact on hepatic glucose and lipid metabolism, offering insights into its potential as a therapeutic target and addressing the need for sex-specific research in liver disease. This study reveals that HKDC1 expression is elevated in obese women with MASH and correlates with liver pathology. In a mouse model, liver-specific HKDC1 knockout (HKDC1^LKO^) protected against Western diet-induced obesity, glucose intolerance, and MASH features, including steatosis, inflammation, and fibrosis. Transcriptomic analysis showed that HKDC1 deletion reduced pro-inflammatory and pro-fibrotic gene expression, while gut microbiome analysis indicated a shift toward MASH-protective bacteria. These findings suggest that HKDC1 may exacerbate MASH progression through its role in metabolic and inflammatory pathways, making it a potential therapeutic target.

## Introduction

Metabolic dysfunction-associated steatohepatitis (MASH), previously referred to as nonalcoholic steatohepatitis (NASH), is a severe and progressive liver disease that affects millions worldwide^1^. It is part of the broader spectrum of metabolic-associated steatotic liver disease (MASLD), a condition characterized by liver fat accumulation, inflammation, and varying degrees of fibrosis in the absence of significant alcohol consumption^2,3^. The global prevalence of MASH has been rising sharply, closely mirroring the increasing rates of obesity and metabolic syndrome, both of which are strongly influenced by sedentary lifestyles and the widespread consumption of Western diets rich in fat and sugar^2,4,5^. Current estimates suggest that up to 25% of the global population may be affected by MASLD, and a significant proportion of these individuals progress to MASH, which carries a higher risk of cirrhosis, hepatocellular carcinoma (HCC), and liver-related mortality^6,7^. Despite its growing burden, therapeutic options for MASH remain limited, highlighting an urgent need for deeper mechanistic understanding and novel therapeutic targets.

While MASH tends to initially affect younger males more frequently, it’s important to recognize that females over 60 experience a higher prevalence of MASH^8^. Reflecting on findings from Japan, it’s particularly concerning to see higher instances of cirrhotic MASH in women (57%) compared to men (43%)^8,9^. It’s also quite alarming that studies indicate women aged 50 and above with MASLD may have a 1.2 times higher likelihood of developing MASH and advancing to severe fibrosis compared to their male counterparts of the same age group^8,9^. A report from the United States highlights a troubling trend between 1999 and 2022, showing an increase in the age-adjusted MASLD-related mortality rate (AAMR) from 0.2 to 1.7 per 100,000^10^. The rise in mortality rates for females, from 0.2 to 2 per 100,000, with an average annual percent change (AAPC) of 11.7% (p<0.001), compared to the rise in males, from 0.2 to 1.3 per 100,000 with an AAPC of 9.3% (p<0.001), underscores the severe impact of this disease on women^10–12^. These sex-specific differences in MASH susceptibility remain poorly understood, and this data compels us to acknowledge that although MASLD may present itself earlier in males, the progression and impact of the disease can be significantly harsher in females.

Central to the development of MASH is the dysregulation of both glucose and lipid metabolism, which significantly alters hepatic energy homeostasis. Excessive accumulation of triglycerides in hepatocytes, driven by increased de novo lipogenesis and impaired fatty acid oxidation, is a hallmark of early MASH. Concurrently, a key feature of metabolic syndrome, insulin resistance exacerbates hepatic glucose production, further disrupting glucose homeostasis. The interplay between these metabolic pathways creates a pro-inflammatory environment within the liver, triggering hepatocellular injury, inflammation, and subsequent fibrogenesis. Understanding the molecular regulators of these metabolic disturbances is critical for identifying new therapeutic targets that could intervene in the early stages of disease progression.

In recent years, hexokinase domain-containing protein 1 (HKDC1) has emerged as a potential player in the metabolic dysfunction observed in liver diseases^13–28^. Unlike other members of the hexokinase family, which are ubiquitously expressed and primarily function in glycolysis, HKDC1 displays unique expression patterns restricted to pathological conditions. Under normal physiological conditions, HKDC1 expression in the liver is minimal. However, its expression is significantly upregulated in response to metabolic stressors such as obesity, insulin resistance, and liver injury, including during the progression of MASLD and MASH. This disease-specific expression pattern suggests that HKDC1 may be specialized in hepatic metabolic reprogramming, particularly in glucose and lipid dysregulation. HKDC1’s involvement in metabolic pathways linked to glucose phosphorylation and lipid metabolism further underscores its potential significance in MASH pathogenesis.

Despite these intriguing findings, the precise role of HKDC1 in MASH, particularly in response to a Western diet, remains largely unexplored. Furthermore, the potential sex-specific effects of HKDC1 in liver disease have not been adequately investigated. Given the heightened risk of MASH progression in postmenopausal women, exploring the role of HKDC1 in female-specific models is critical for understanding how this enzyme contributes to sex-based disparities in disease outcomes.

In this study, we aim to elucidate the role of HKDC1 in the development of Western diet-induced MASH, focusing specifically on female mice. Our findings provide novel insights into the metabolic underpinnings of MASH and address the critical need for research on sex-specific mechanisms in liver disease.

## Materials and Methods

### Animal Experiments

All mice were housed in a temperature- and humidity-controlled specific pathogen-free barrier facility with ad libitum access to food (Envigo-7912, IN, USA), water, and 12 hours of dark and light cycles. All procedures were approved by The University of Illinois Chicago Animal Care Committee. HKDC1^fl/fl^ mice^26,28,29^ were crossed with Alb-Cre mice (Jackson Laboratory, Strain:003574, ME, USA) to generate Alb-Cre^+/-^ HKDC1^fl/fl^ and HKDC1^fl/fl^ mice as before^30^. In this study, only female mice were used. When mice reached 8 weeks of age, Western diet (Research diet, D12079Bi, NJ, USA) or normal chow (Envigo-7912, IN, USA) were given for 28 weeks.

### Body composition, glucose homeostasis, and metabolic rate

Whole body fat, lean, and fluid mass were measured with a minispec LF50 Body Composition Analyzer (Bruker, Billerica, MA)^31,32^. Glucose (2mg glucose ip/g) and pyruvate (2mg sodium pyruvate ip/g) tolerance tests were performed in overnight fasted mice. Blood glucose was measured from a lateral tail vein with a glucometer (Accu-check, Roche). Energy expenditure, VO2, VCO2, respiratory exchange ratio, food intake, and activity were measured using Promethion Systems (Sable Systems International, Las Vegas, NV)^33^.

### Study design and participants

We included 95 women with obesity who underwent bariatric surgery at the Virgen de la Arrixaca University Hospital (Murcia, Spain) between January 2020 and December 2021. Inclusion criteria included a signed informed consent, age 18 to 65 years, body mass index (BMI) ≥ 35 kg/m^2^ or ≥ 30 kg/m^2^ with significant obesity-related comorbidities, and duration of obesity ≥ 5 years. Exclusion criteria were evidence of liver disease other than MASLD (including viral hepatitis, medication-related disorders, autoimmune disease, hepatocellular carcinoma, hemochromatosis, Wilson’s disease, and familial/genetic causes), previous history of excessive alcohol use, treatment with any drugs potentially causing steatosis such as tamoxifen, amiodarone, and valproic acid, or subjects who declined to participate. Study groups were defined based on the histological evaluation of liver biopsies according to the SAF (Steatosis, Activity, Fibrosis) classification system as (1) patients without MASLD (no-MASLD), i.e. liver without any histological alteration; (2) patients with metabolic dysfunction-associated steatotic liver (MASL), defined by the presence of at least grade 1 (5%) liver steatosis, with or without ballooning or lobular inflammation but not both at the same time; and (3) patients with MASH, defined by the presence of at least grade 1 steatosis, ballooning, and lobular inflammation, with or without fibrosis ^34^. The study was performed per the Declaration of Helsinki according to local and national laws and was approved by the Ethics and Clinical Research Committees of the Virgen de la Arrixaca University Hospital (ref. number 2020-2-4-HCUVA).

#### Anthropometric and biochemical measurements

The body mass index (BMI) of the individuals was calculated by Quetelet’s formula: *BMI = weight in kg/(height in meters)*^2^. Preoperative clinical data were collected on the day of the surgery after an overnight fast of at least 12 hours and serum was separated by centrifugation. Laboratory measurements including glucose, alanine aminotransferase (ALT), aspartate aminotransferase (AST), and gamma-glutamyltransferase (GGT) levels were performed using the Cobas Analyzer c702 (Roche) following standardized methods. The levels of glycated hemoglobin (HbA1c) were measured in blood with the glycohemoglobin analyzer HLC®-723G8 (Tosoh Bioscience). Insulin levels were measured using the Cobas Analyzer e801 (Roche). Insulin resistance was determined by the Homeostasis Model Assessment of Insulin Resistance (HOMA-IR) index and was calculated as insulin (µU/mL) x glucose (mmol/L)/22.5^35^.

### Food Intake

Mice were placed in individual and mouse cages (Research Diets Inc., New Brunswick, NJ, Canada) to measure individual food intake^36^. An acclimatation period was carried out to limit the stress of the mice associated with this new environment. By this BioDaq system, the mouse feeders are connected to a computer, which very accurately calculates the amount of food consumed by each mouse over 24 hours.

### Fecal Bile acid and NEFAs

Feces were collected from mice over a 24-hour period using wire-bottom cages. Bile acid was extracted using the method of Humbert et al.,^37^ Briefly, 2mL of 0.1M NaOH was added to 100mg of feces for one hour at 60C. Then, 4mL of miliQ water was added, and the mixture was homogenized at maximum speed. The homogenized sample was centrifuged at 20000g for 20 minutes. Mouse total bile acids assay kit (Crystal Chem) was used to quantify the fecal bile acid.

Fecal NEFAs were extracted using a modified Folch method^38^. Briefly, approximately 100 mg of fecal sample was homogenized in 1 mL of chloroform:methanol mixture (2:1, v/v). The homogenate was vortexed vigorously for 5 minutes and centrifuged at 3,000 × g for 10 minutes at 4°C, and the lower organic phase, which contains the lipids, was carefully collected. The collected organic phase was then evaporated under a gentle stream of nitrogen to dryness. The dried lipid residue was reconstituted in isopropanol and subsequently analyzed for non-esterified fatty acids (NEFAs) using the Wako NEFA estimation kit as before^32^.

### Histology and pathology assessment

Formalin-fixed livers were processed by the Research Histology and Tissue Imaging Core of the University of Illinois at Chicago, and stained with Hematoxylin & Eosin or picrosirius red/fast green^33^. Pathological features of the liver sections were graded in a blinded fashion, following Kleiner’s scoring system^39^.

### Human liver organoid (HLO) isolation, culture, and treatment

Cryopreserved human liver samples were thawed at 37 °C and placed in a petri dish containing ice-cold 15 mL of DMEM/F-12 with 15 mM HEPES and 10% FBS. Samples were disassociated using a solution containing DMEM/F-12 with 15 mM HEPES, DNase I (Gibco), and collagenase type IV (Gibco) and incubated at 37 °C for 15 min. Then, samples were resuspended and left for 1 min to allow bigger pieces to precipitate, and the supernatant was transferred to a new tube in ice. This step was repeated 4 times. Once the tissue was completely disaggregated, samples were centrifuged, the supernatant was discarded, and the pellet was resuspended in 10 mL of TrypLE (Gibco) and incubated for 20 min at 37 °C. After this incubation, samples were resuspended, and 10 mL of Advanced DMEM with DNase I was added. Samples were centrifuged, and the pellet was resuspended in cold Ammonium Chloride Solution (StemCell Technologies) and Advanced DMEM with DNase I and incubated for 5 min in ice. Then, samples were centrifuged, the pellets were washed with DMEM/F-12 with 15 mM HEPES and 10% FBS, centrifuged again, and resuspended in 2 mL of Advanced DMEM + DNase I. The number of viable cells was determined using a Trypan blue solution (Sigma-Aldrich). To initiate the cultures, after centrifuging, the pellet was resuspended in the volume of Matrigel necessary to obtain a density of 20,000 live cells per dome. A total of 40 µL of Matrigel-containing cells were placed in each well, and the plate was incubated at 37 °C for 30 min before adding 500 µL of HepatiCult Organoid Initiation Media (StemCell Technologies). The medium was changed every 2-3 days until the organoid cultures were ready for passaging (∼ 14 days). Before performing the experiments, organoid samples were passaged and maintained for at least two passages in HepatiCult Organoid Growth Media (StemCell Tecnologies). Then, organoids were collected and dissociated into small fragments (30 -100 μm). The number of fragments in suspension was determined using a light microscope. The volume required to obtain 1,000 fragments/well was calculated, transferred to a new tube, centrifuged, and the pellet was resuspended in the pertinent amount of Matrigel. For viability assays, 8 µL/well of Matrigel-containing fragments were placed in 96-well white opaque plates (Greiner Bio-One, Kremsmünster, Austria). For RT-qPCR, 20 µL/well was placed in 48-well plates. The plates were incubated at 37 °C for 30 min before adding HepatiCult Organoid Growth Media and maintained for 5 days, changing the media after 2 days. On day 5, the medium was changed to HepatiCult Organoid Differentiation Media (StemCell Technologies) for 10 days before treating the cells with 400 µM or 600 µM of PA. Organoid cultures were incubated for 5 days, and the different treatments with media change were performed every 2-3 days. To extract RNA from HLO, organoids were first isolated from Matrigel using an ice-cold Cell Recovery Solution (Corning). The solution was added directly into the wells and the plates were placed in ice for 30 min under agitation. After this, samples were collected in 1.5 mL nuclease-free tubes and centrifuged at 2,000 x g for 5 min at 4 °C. Samples were then washed 4 times with ice-cold PBS. RNA extraction was performed using the PicoPure RNA Isolation Kit (Applied Biosystem, Waltham, MA, USA) following the manufacturer’s instructions.

### Glucose and Insulin tolerance test

Mice were fasted for 16h for GTT and 4h for ITT. Mice were injected ip with 2mg/kg glucose (for GTT) or 0.5 Units/kg bwt of Humalog insulin (for ITT; EliLily&Co, Indianapolis, IN, USA). Blood glucose levels were determined similarly to the aforementioned glucose tolerance tests.

### Liver biopsies collection and sample processing

Liver biopsies of at least 1 cm^2^ were obtained from patients on the day of the surgery. The biopsy specimens were collected in 1.5 mL nuclease-free tubes containing 1 mL of RNA stabilization Solution (RNAlater, Sigma) for RT-qPCR analyses, maintained overnight at 4°C and kept at - 80°C until use.

### Metabolic endpoints measured in plasma

Mice were sacrificed by decapitation 4h after food withdrawal at 0700am, and trunk blood was collected to determine levels of NEFA, triglyceride (TG), cholesterol (Wako Diagnostics, Richmond VA), and ALT (Pointe Scientific, Canton, MI). To assess hepatic TG content, hepatic lipids were extracted from frozen livers in isopropanol, and TG was measured as previously published^26,28^.

### RNAseq and qPCR

RNA was extracted using Trizol Reagent (Life Technologies, Carlsbad, CA) and used to perform RNA-seq in an Illumina Platform PE150 with bioinformatics analysis (Novogene, Inc, Sacramento, CA) or to perform qPCR as previously described^30,40,41^. Data has been uploaded to NCBI Geo (Accession Number GSE283058).

### 16S rRNA Gene Sequencing from Mouse Cecal Matter

Cecal contents were collected from mice and immediately flash-frozen in liquid nitrogen before storage at – 80°C. Genomic DNA was extracted using the Qiagen DNeasy PowerSoil Kit following the manufacturer’s protocol to ensure high-quality microbial DNA recovery. The V4 hypervariable region of the 16S rRNA gene was amplified by Novogene Inc. using standard in-house protocols. Raw sequence data were processed and analyzed using QIIME2, with taxonomic assignment performed against a reference database. The raw data can be accessed via sequence accession PRJNA1191474

### Statistical analysis

Values are represented as means ± standard error of deviation (SD). The normal distribution of the study variables was assessed by the Kolmogorov– Smirnov test. The categorical data analysis was performed by the χ2 test for intragroup differences and the Pearson χ2 test for intergroup differences. Relationships between pairs of continuous variables were determined based on the Spearman correlation test. Data were analyzed by *t*-test, one-way or 2-way ANOVA followed by a Tukey or Bonferroni post-hoc test when applicable. Analysis of RNAseq data and enrichment analysis of DEG was performed by Novogene, Inc. Differentially regulated metabolites, and enrichment analysis of metabolomics was performed with Metware Inc. The statistical analyses were performed using GraphPad Prism 8 (GraphPad Software, La Jolla, CA). p-values less than 0.05 were considered significant.

## Results

### Hepatic HKDC1 gene expression is upregulated in obese women with MASH

We have previously shown a positive correlation between HKDC1 protein levels and features of MASLD in humans^42^ and mice on a choline-deficient amino acid-defined diet^30^. This observation piqued our interest in studying the role of HKDC1 overexpression in MASLD progression, particularly in females, because HKDC1 was first identified as a risk factor in maternal hyperglycemia^43,44^. To gather further evidence, we collected liver samples from obese women who had undergone bariatric surgery [cases with steatotic (SLD) or MASH diagnosis]. Quantitative PCR analysis revealed that although there was no significant change in the mRNA levels of *HK1, HK2,* or *GCK* (Fig 1 A-C), there was a significant increase in *HKDC1* gene expression in obese women with MASH compared to that in healthy livers (Fig 1D). Unsurprisingly, in these patients, a strong positive correlation was observed between pathological scoring of liver biopsies, including steatosis, inflammation, ballooning, fibrosis, SAF score, and stages of MASLD. In addition, CK18, a marker of liver injury, was significantly correlated with hepatic HKDC1 mRNA levels (Table 1).

**Fig 1:**
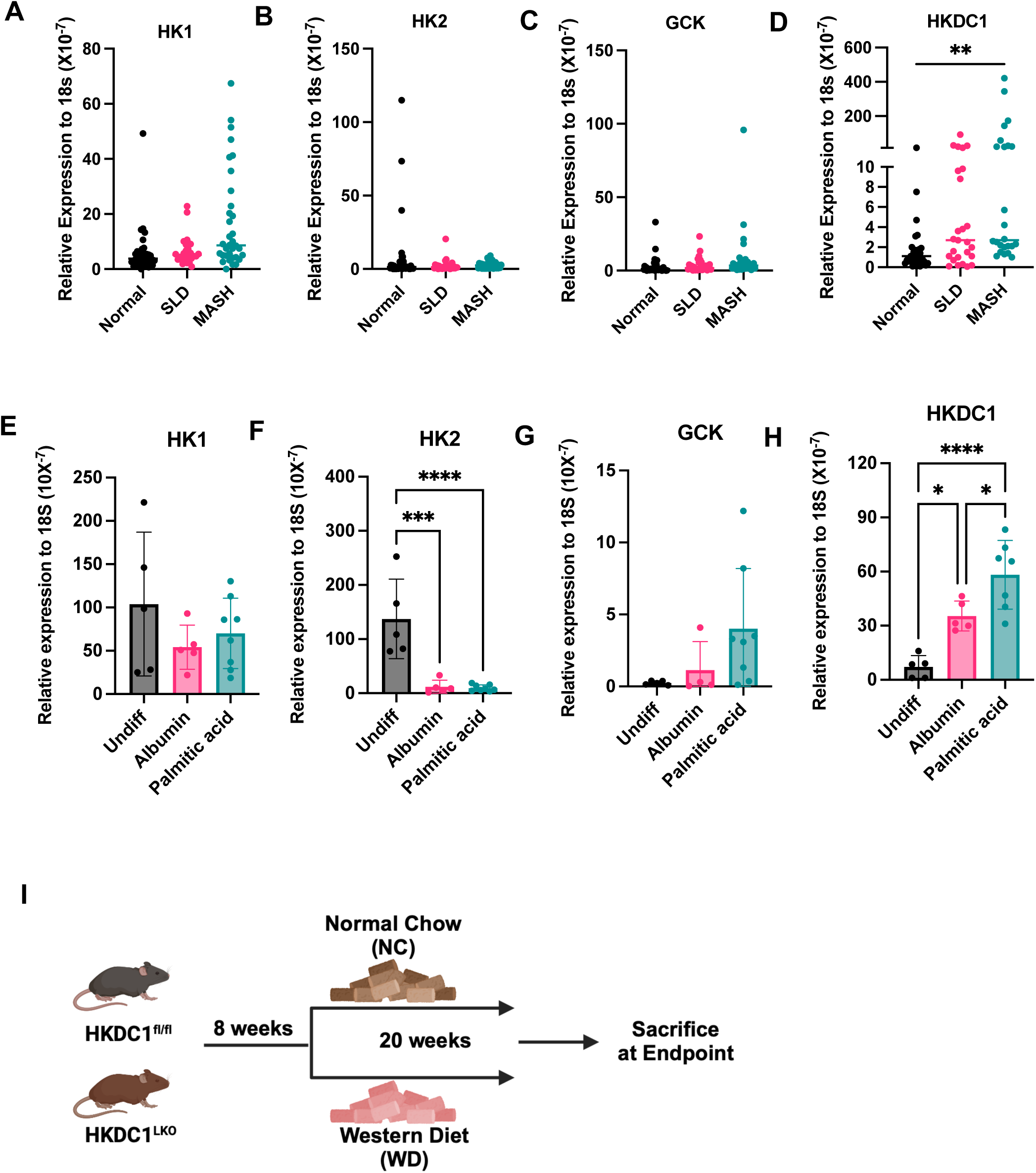
HKDC1 gene expression is upregulated in the liver of obese women with MASH. (**A-D**) HK1, HK2, GCK, and HKDC1 expression levels in normal, steatotic, and steatohepatitic livers from women with obesity (n = 24-32) (**E-F**) HK1, HK2, GCK, and HKDC1 expression levels in undifferentiated, Differentiated - treated with albumin or palmitic acid (n = 5-7). (**I**) Study design of HKDC1 hepatic knockout and WT littermates fed with normal chow (NC) or western diet (WD). Data were analyzed by ordinary one-way ANOVA with Dunnett correction, *p < 0.05, **p < 0.01, ***p < 0.001, ****p < 0.0001.

**Table 1:**
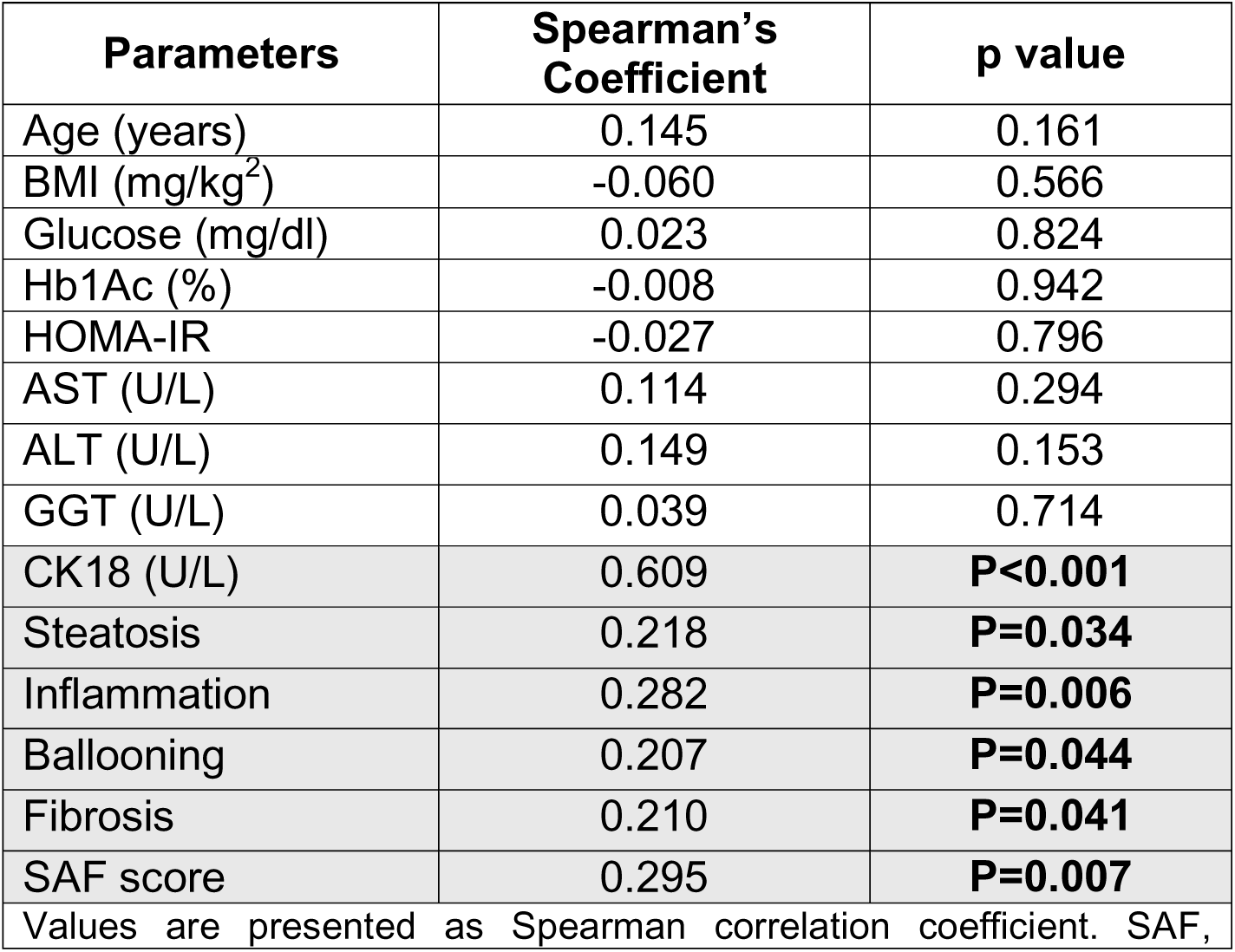
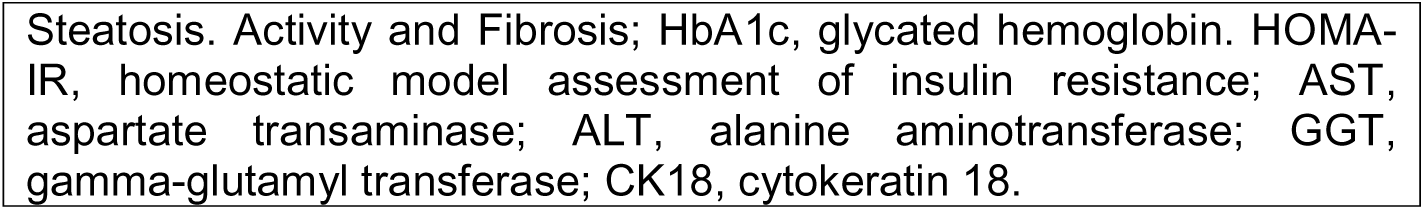
Correlations between hepatic HKDC1 expression and histological and biochemical parameters related to MASLD.

| Parameters | Spearman's Coefficient | p value |
| --- | --- | --- |
| Age (years) | 0.145 | 0.161 |
| BMI (mg/kg<sup>2</sup>) | -0.060 | 0.566 |
| Glucose (mg/dl) | 0.023 | 0.824 |
| Hb1Ac (%) | -0.008 | 0.942 |
| HOMA-IR | -0.027 | 0.796 |
| AST (U/L) | 0.114 | 0.294 |
| ALT (U/L) | 0.149 | 0.153 |
| GGT (U/L) | 0.039 | 0.714 |
| CK18 (U/L) | 0.609 | **P<0.001** |
| Steatosis | 0.218 | **P=0.034** |
| Inflammation | 0.282 | **P=0.006** |
| Ballooning | 0.207 | **P=0.044** |
| Fibrosis | 0.210 | **P=0.041** |
| SAF score | 0.295 | **P=0.007** |

Values are presented as Spearman correlation coefficient. SAF,
Steatosis. Activity and Fibrosis; HbA1c, glycated hemoglobin. HOMA-IR, homeostatic model assessment of insulin resistance; AST, aspartate transaminase; ALT, alanine aminotransferase; GGT, gamma-glutamyl transferase; CK18, cytokeratin 18.

Further, we used cryopreserved human liver samples to form *ex vivo* liver organoids, which were differentiated and then treated with 400uM palmitic acid (PA) to mimic a high lipid environment such as seen during MASH^35^. Our data shows that while there was no change in mRNA levels of *HK1* and *GCK* (Fig 1 E, G), *HK2* levels were significantly downregulated in differentiated liver organoids both untreated and treated with PA compared to undifferentiated HL (Fig 1 F). More importantly, HKDC1 levels were not only significantly upregulated in differentiated HLO compared to undifferentiated HLO but were also increased after PA treatment (Fig 1H), suggesting that a high lipid environment can regulate HKDC1 expression.

To study the role of HKDC1 in the pathogenesis of MASH, we used HKDC1 floxed (HKDC1^fl/fl^) mice^30,32,42^ and Albumin Cre (Jackson Labs) mice to generate liver-specific HKDC1 knockout (HKDC1^LKO^). At 8 weeks of age, HKDC1^LKO^ and HKDC1^fl/fl^ (Controls) female mice were fed either normal chow (NC) or Western diet (WD) for 20 weeks (Fig 1I). Mice fed with NC showed the role of hepatic HKDC1 under normal physiological conditions and served as the baseline. The WD challenge mimics MASH pathogenesis, which allows exploration of the role of HKDC1 in the pathogenesis of MASH.

### Hepatic HKDC1 deletion protects mice from WD-induced obesity

During the 20 weeks of diet intervention, we measured body weight and body composition, which was examined using nuclear magnetic resonance (NMR) every 4 weeks. In the NC-fed group, we did not observe any difference in body weight, fat mass, lean mass, and free body fluid percentage between the groups (Fig 2A-F). However, in the mice fed WD, we noticed significantly less weight gain in female HKDC1^LKO^ mice starting from 16 weeks of age and further expanded throughout the rest of the study period (Fig 2A). This less weight gain in WD-fed female HKDC1^LKO^ mice was due to significantly lower fat mass (Fig 2B, E) concomitant with modestly preserved lean mass compared to HKDC1^fl/fl^ females (Fig 2C, F). No difference was noted in the free-body fluid between the two groups (Fig 2D). Thus, HKDC1 hepatic ablation protects female mice from WD-induced obesity by reducing fat mass gain.

**Fig 2:**
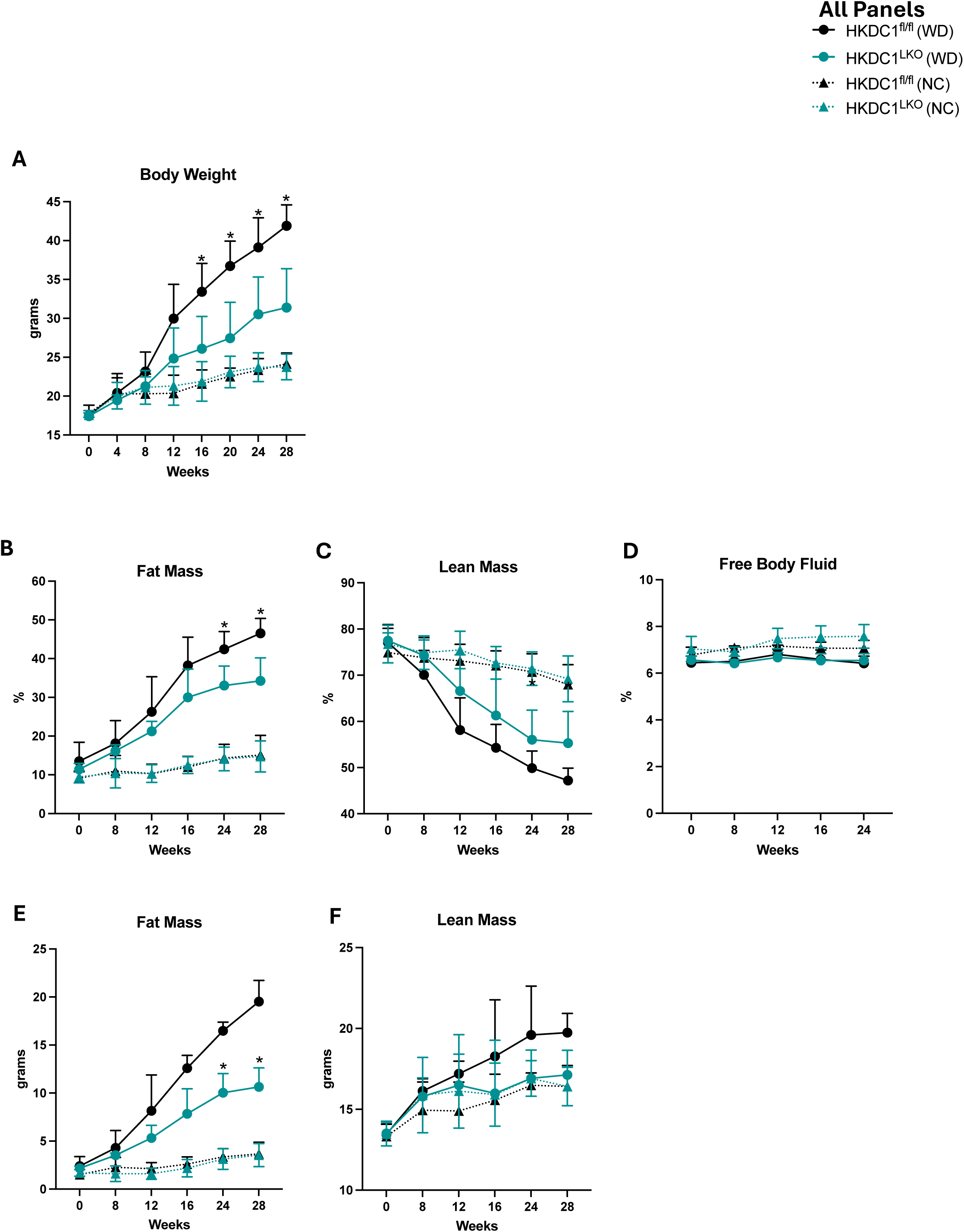
Hepatic HKDC1 deletion protects mice from WD-induced obesity. Female HKDC1^fl/fl^ and HKDC1^LKO^ were subject to body weight (A) and NMR body composition measurements, including fat mass (B), lean mass (C), and free body fluid (D) at indicated times. Absolute values for (E) fat mass and (F) lean mass. N = 4-5 for each genotype per condition. Data were analyzed by two-way ANOVA with Šidák correction, *p < 0.05.

### Hepatic HKDC1 ablation does not affect overall energy expenditure

To understand the reason for the lower body weight and fat mass in female HKDC1^LKO^ mice under WD feeding, the same mice were subjected to indirect calorimetry in the metabolic cage at 24 weeks (at 16 weeks under diet treatment). No significant difference was observed in the energy expenditure (Fig 3A-C), oxygen consumption (VO_2;_ Fig 3D-F), CO_2_ production (VCO_2_; Fig 3G-I), and respiratory exchange ratio (RER; Fig 3J-K) between the female HKDC1^LKO^ mice and HKDC1^fl/fl^ mice fed WD on individual days and nights and the average reading of days and nights. Locomotion and food intake also did not change significantly (Fig 3L and 3M). In NC-fed mice, there was no difference between female HKDC1^LKO^ mice and HKDC1^fl/fl^ mice in VO_2_, VCO_2_, RER, or food intake (S Fig 1A-D). To confirm our food intake data in the WD group, we also used BioDaq to measure food intake more accurately, and our data confirms that HKDC1^LKO^ mice do not eat less than the control mice at the start (S Fig 1E) and towards the end of the study (S Fig 1F). Overall, the knockout of HKDC1 in hepatocytes does not alter the energy balance in female HKDC1^LKO^ compared to HKDC1^fl/fl^ mice.

**Fig 3:**
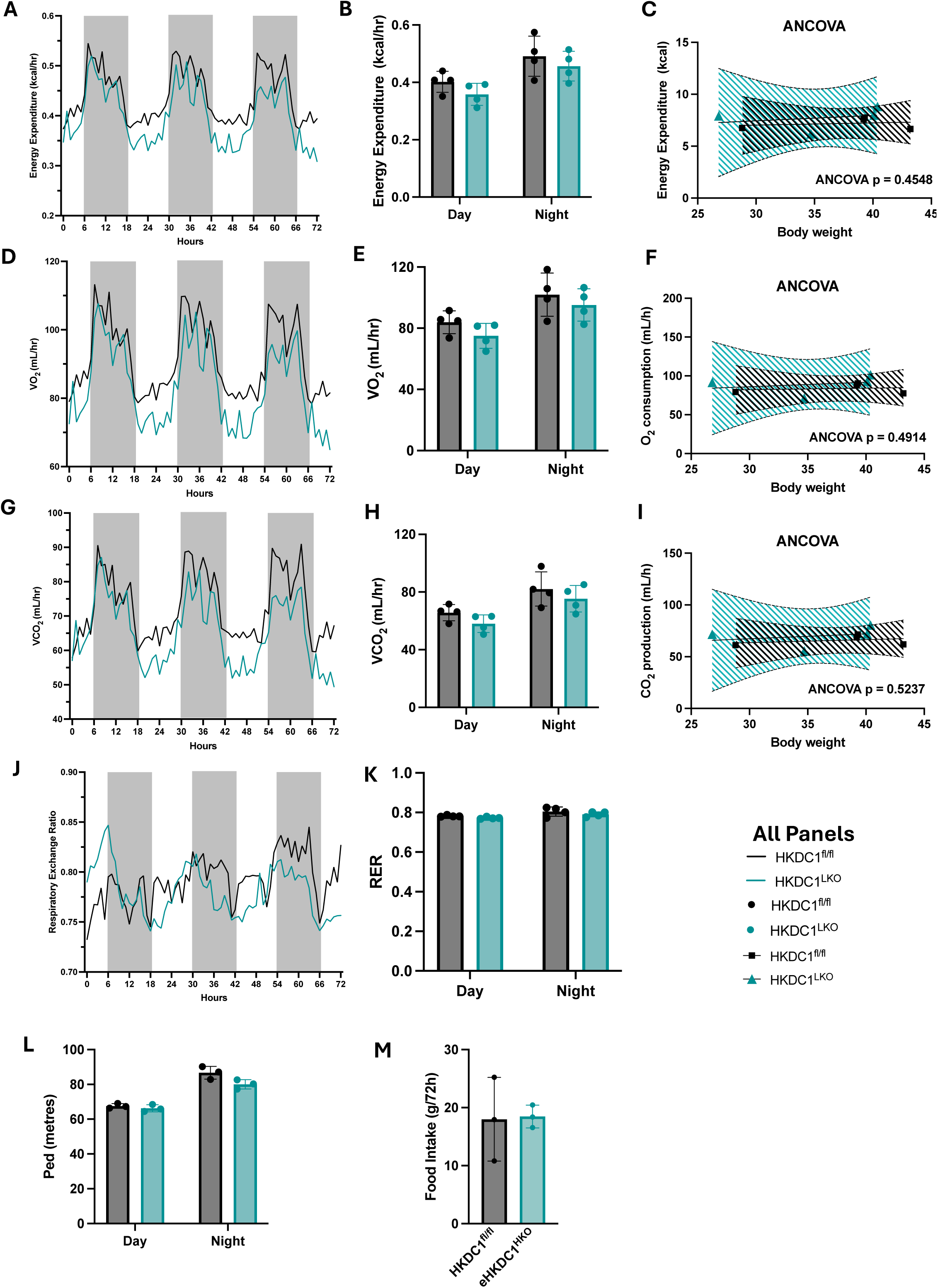
Hepatic HKDC1 ablation does not affect the overall energy balance in WD-fed conditions. At 20 weeks, female HKDC1^fl/fl^ and HKDC1^LKO^ fed with WD were subjected to indirect calorimetry and food intake measurement. Individual day and night energy expenditure (**A-C**) Energy expenditure, (**D-F**) O_2_ consumption, (**G-I**) CO_2_ production, (**J-K**) respiratory exchange ratio, (**L)** locomotion, and (**M**) cumulative food intake (n = 4). Data were analyzed using two-way ANOVA with Dunnett correction.

### Hepatic HKDC1 leads to glucose intolerance and insulin resistance only under the WD-fed condition

WD affects glucose homeostasis by increasing glucose intolerance and insulin resistance. To examine how these are impacted by hepatic HKDC1, we conducted intraperitoneal glucose tolerance and insulin tolerance tests in female HKDC1^LKO^ and HKDC1^fl/fl^ mice prior to diet challenge (8 weeks of age) and 16 weeks (24 weeks of age) into diet feeding. Prior to the diet challenge, we did not observe any difference in glucose tolerance between female HKDC1^LKO^ and HKDC1^fl/fl^ mice (Fig 4A-B). While mice fed NC for 16 weeks did not exhibit any difference in glucose tolerance (Fig 4C-D), on the WD, HKDC1^LKO^ mice showed better glucose tolerance than HKDC1^fl/fl^ mice (Fig 4C-D). No significant differences were observed in the monthly random blood glucose levels (Fig 4E). However, HKDC1^LKO^ had lower fasting blood glucose than controls (Fig 4F). Further, HKDC1^LKO^ mice on the WD were more sensitive to insulin in the insulin tolerance test than HKDC1^fl/fl^ mice (Fig 4G), and as expected, no changes were observed among the groups on the NC (Fig 4G).

**Fig 4:**
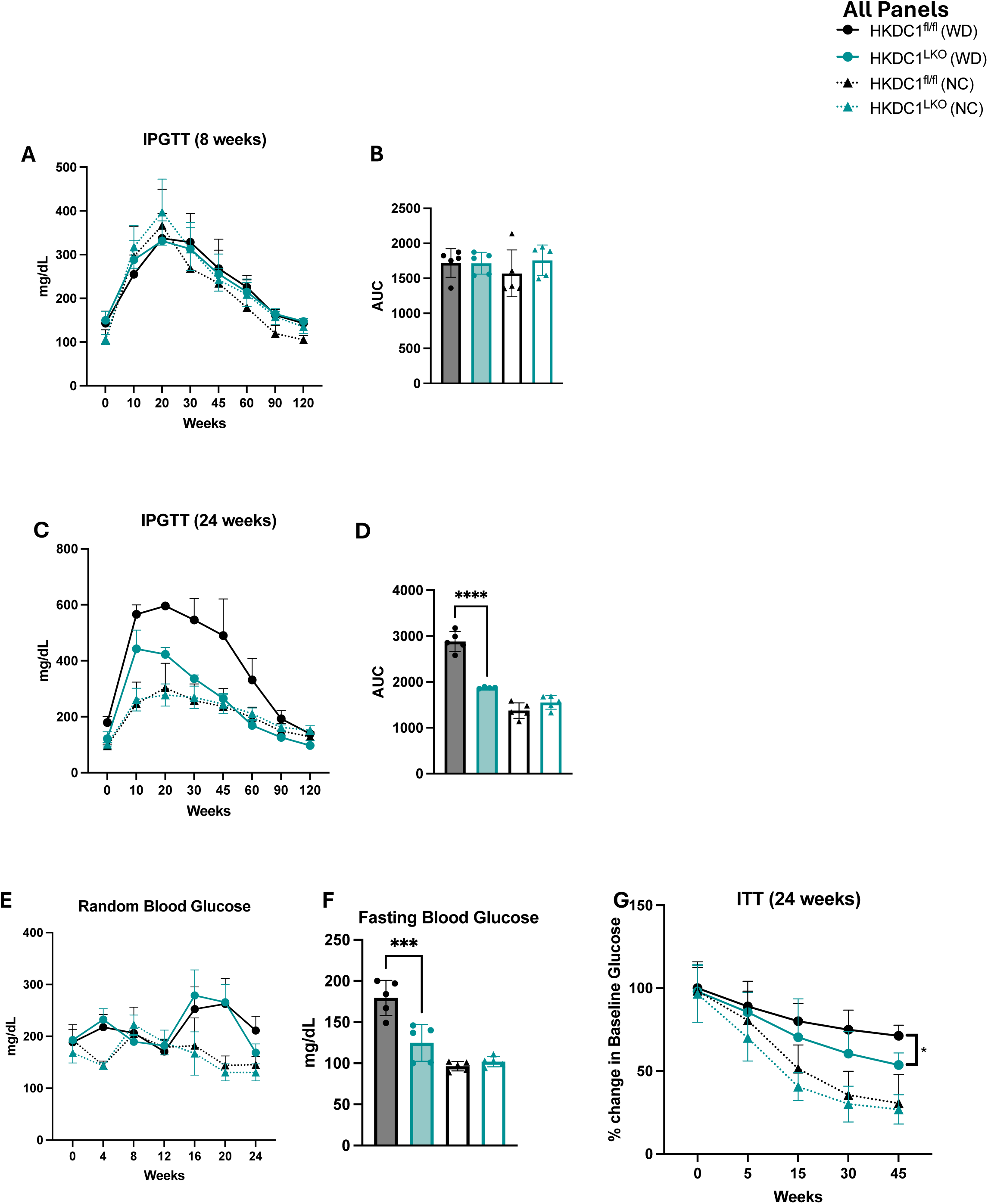
Hepatic HKDC1 leads to glucose intolerance and insulin resistance only under the WD-fed condition. Female HKDC1^fl/fl^ and HKDC1^LKO^ mice were subjected to intraperitoneal glucose tolerance tests (IPGTT) at (**A**) before the diet challenge at 8 weeks and (**C**) **at** 24 weeks. Their respective area under curves are shown in B and D. (**E**) Random blood glucose was measured at indicated weeks and (**F**) fasting blood glucose at 24 weeks. (**G**) The insulin tolerance test (ITT) was carried out at 24 weeks. N = 4-5 for each genotype per condition. Data was analyzed by two-way ANOVA with Šidák correction for A, C, E, G and one-way ANOVA with Šidák correction for B, D, F. *p < 0.05, ***p < 0.001, and ****p < 0.0001.

### Liver HKDC1 ablation protects against WD-induced MASH progression

The most prominent features of the liver in MASH patients and mice are hepatomegaly, steatosis, inflammation, and inflammation-induced fibrosis. Interestingly, HKDC1^LKO^ mice had smaller livers than HKDC1^fl/fl^ mice (Fig 5A). Histopathological analysis revealed that HKDC1^LKO^ mice had markedly less steatosis (Fig 5B) and fewer lipid droplets (Fig 5C) than HKDC1^fl/fl^ mice. The NAFLD activity score (NAS) is crucial for evaluating MASLD as it provides a standardized, quantitative assessment of liver histology, scoring key features like steatosis, inflammation, and hepatocellular ballooning. This helps distinguish MASLD from its more severe form, MASH, which is associated with higher risks of fibrosis and liver complications. Liver-specific HKDC1 ablation seems to be hepatoprotective as reflected by the overall lower NAS score (Fig 5D) and including steatosis (Fig 5E), ballooning (Fig 5F), and inflammation (Fig 5G) scores. HKDC1^LKO^ mice also had lower ALT values, supporting the key role of HKDC1 in MASH progression (Fig 5H). Since histomorphology indicated a significant increase in inflammation, cytokine gene expressions, including *TNF*!l!, *Trem*, and *Mcp1,* were also assessed. We noted a significant reduction in the expression of these genes in the livers of HKDC1^LKO^ mice compared to those of HKDC1^fl/fl^ mice (Fig 5I), suggesting that the livers of HKDC1^LKO^ mice had less inflammation and thus likely less damage. To confirm this, we performed Sirius red staining, and indeed, HKDC1^fl/fl^ mice fed a WD diet showed elevated fibrosis in their liver compared to HKDC1^LKO^ mice (Fig 5B), which was also manifested by the fibrosis score (5J). In agreement with the Sirius red staining results, core fibrosis gene expression, including *Col1a* and *Col3a*, was significantly reduced in the livers of female HKDC1^LKO^ mice (Fig 5K). The decrease in whole-body adiposity of HKDC1^LKO^ was also reflected in lower iWAT (Fig 5L) and significantly lower eWAT depot weights (Fig 5M). Histologically, the adipocytes in both groups did not seem to be morphologically different however HKDC1^fl/fl^ mice had more areas with inflammation (Fig 5N). We also quantified adipocytes in the iWAT samples (Fig 5O) and levels of the thermogenic genes CPT 1&2 in iWAT and eWAT (S Fig 2F-G) but found no statistically significant changes.

**Fig 5:**
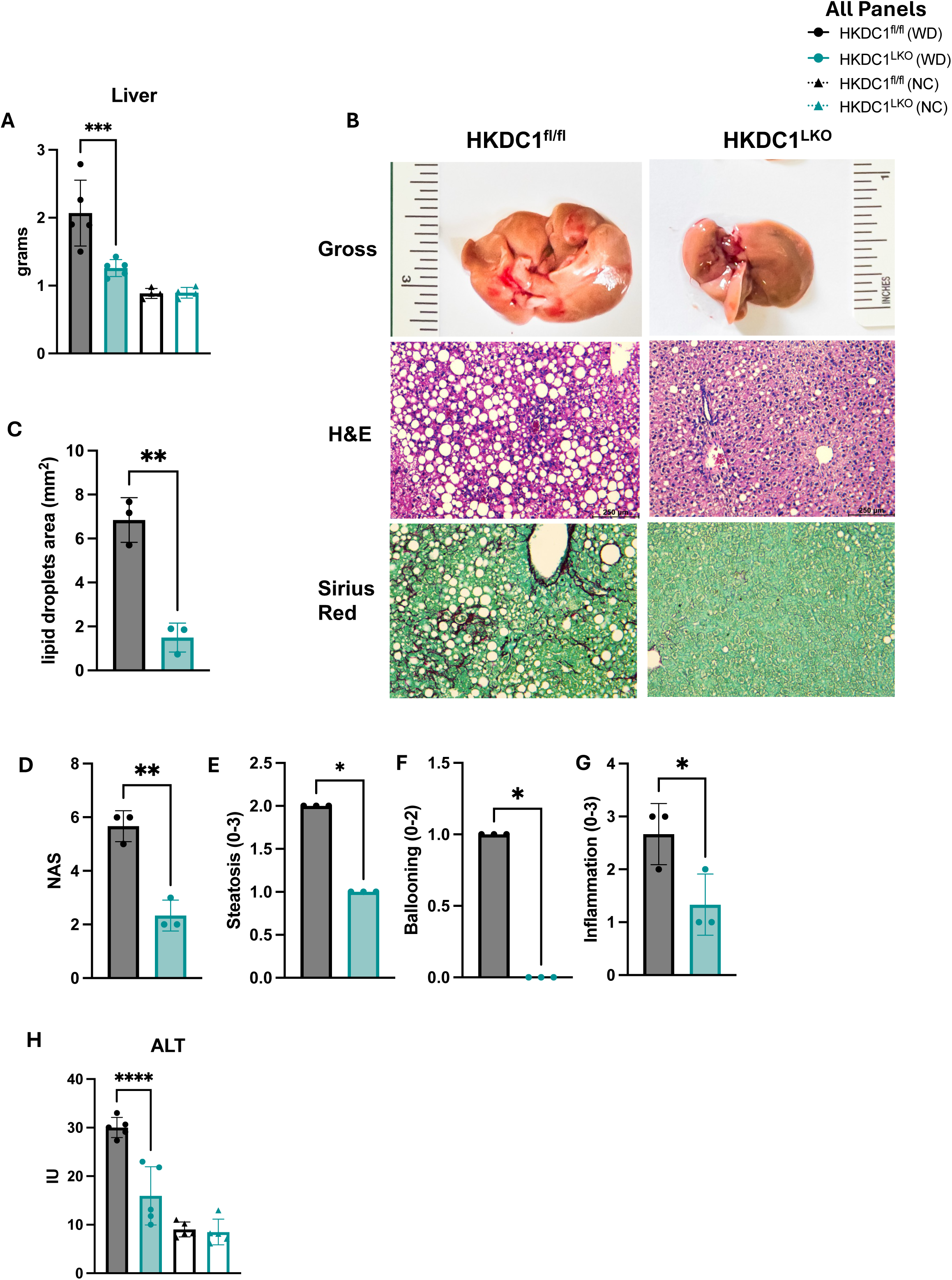
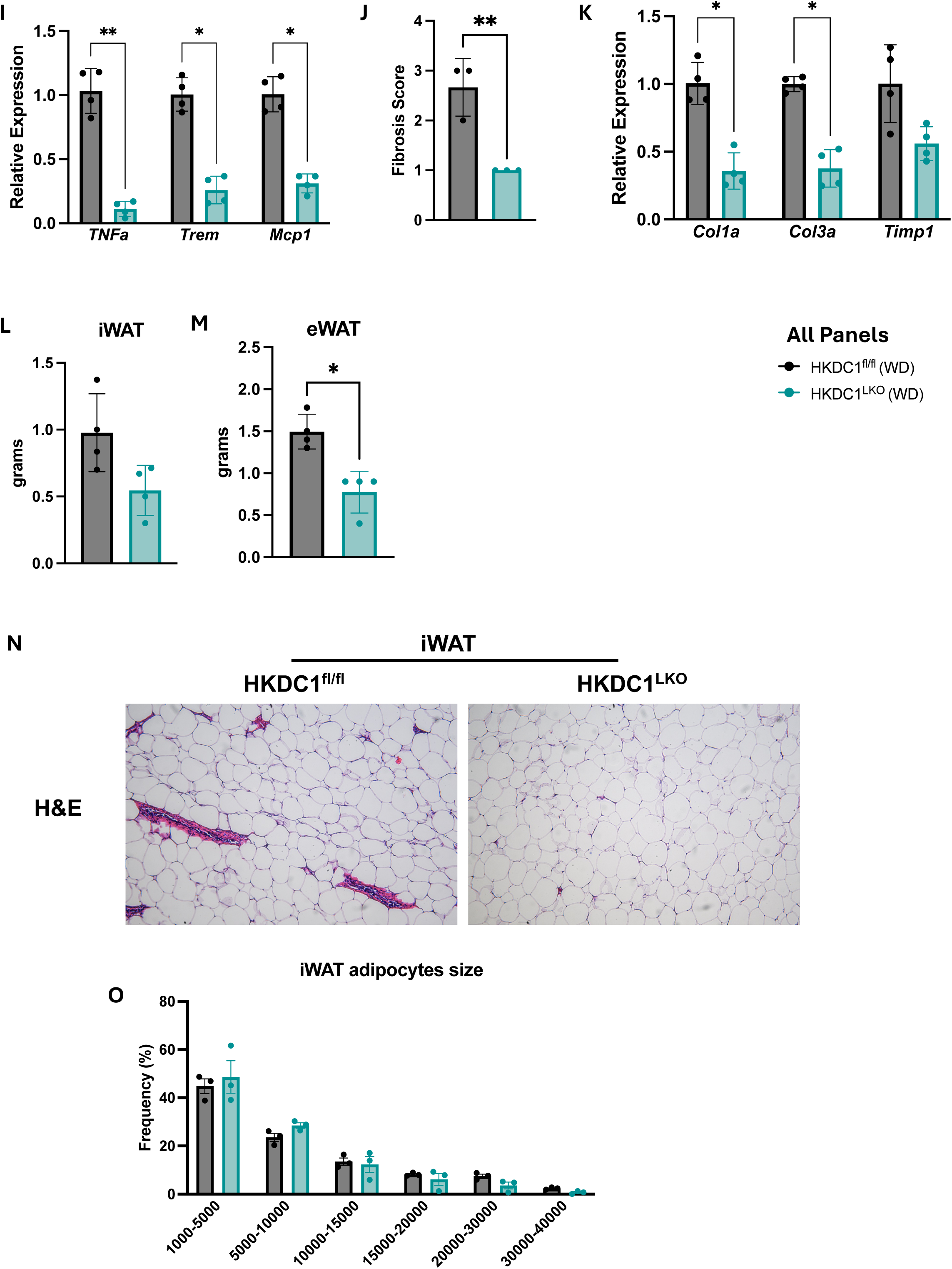
Liver HKDC1 deficiency protects against WD-induced MASH pathogenesis. Female HKDC1^fl/fl^ and HKDC1^LKO^ mice were fed NC or WD for 20 weeks. At the end of the experimental period mice were euthanized and analyzed for (**A**) liver weight, (**B**) histological analysis of liver tissue by H&E and Sirius red, (**C**) lipid droplets area was quantified from H&E images of individual mice using Image J (**D**) NAFLD activity score (NAS) score consisting of (**E**) steatosis, (**F**) ballooning, and (**G**) inflammation. (**H**) serum ALT values from 4h fasted mice. (**I**) mRNA expression of cytokine genes *TNF*!l!, *Trem*, and *Mcp1* expressed as fold change relative to HKDC1^fl/fl^, (**J**) fibrosis score and (**K**) mRNA expression of fibrosis genes *Col1a*, *Col3a*, and *Timp1* expressed as fold change relative to HKDC1^fl/fl^. Adipose depots were also collected and weighed; shown here are the weights of (**L**) inguinal (iWAT) and (**M**) epididymal (eWAT). (**N**) histological analysis of iWAT tissue by H&E. N = 3-5 for each genotype per condition. Data were analyzed by *t*-test for C-G, J, L, M and one-way ANOVA with Dunnett correction for I, K, two-way ANOVA with Šidák correction for A and H. *p < 0.05, **p < 0.01, ***p < 0.001, and ****p < 0.0001.

### Liver HKDC1 ablation acts as a metabolic reset button in MASH

In accordance with histomorphology, serum and liver triglyceride (TG) levels were lower in HKDC1^LKO^ mice (Fig 6A-B). We did not find any significant change in circulatory NEFAs in four-hour fasted mice (S Fig 2A), and qPCR analysis of some important mediators of fatty acid metabolism did not reveal any significant changes except for *Cyp4A10* that showed marked induction in HKDC1^LKO^ livers (S Fig 2B). Cholesterol plays a key role in MASLD as its accumulation in the liver contributes to metabolic dysfunction, inflammation, and cellular stress, all driving disease progression. High cholesterol levels exacerbate steatosis and can lead to the development of MASH. Supporting our data that HKDC1 ablation is hepatoprotective against diet-induced MASH, serum (Fig 6C-E) and hepatic cholesterol (Fig 6F) levels were also reduced in HKDC1^LKO^ mice. Further, the expression of important cholesterol metabolism genes, including *Hmgcr, Ldlr,* and *Fabp1,* showed an opposite trend to the dysregulation seen in MASLD (Fig 6G). We did not notice any difference between HKDC1^LKO^ and HKDC1^fl/fl^ mice in the NC condition in liver TG and cholesterol, serum ALT, TG, NEFA, and cholesterol levels (Fig 6A-F; S Fig 2A). Thus, these findings suggest that only under WD challenge HKDC1 is a MASH pathogenesis contributor, which increases steatosis, inflammation, and fibrosis of the liver, and that a possible mechanism by which HKDC1 upregulation exacerbates MASH development is through dysregulating cholesterol metabolism.

**Figure 6:**
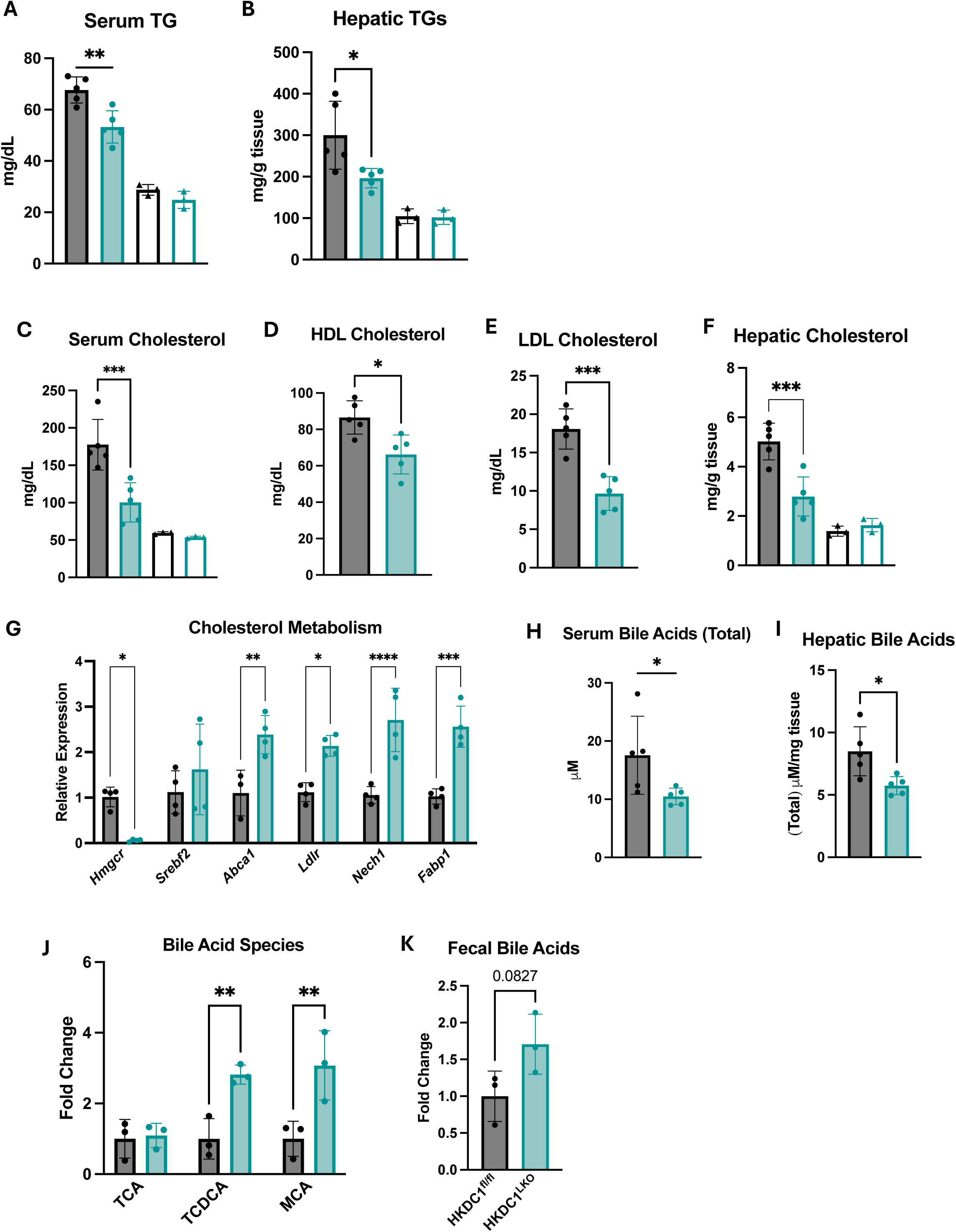
HKDC1 deletion normalizes lipid and cholesterol metabolism in MASH diet-fed mice. At the terminal endpoint. Mice were fasted for 4h and then sacrificed and livers were harvested. Blood was collected, and serum was separated by centrifugation. Serum and liver homogenates were used to assess the levels of (**A**) serum and (**B**) hepatic triglycerides (TGs), (**C**) serum, (**D**) HDL and (**E**) LDL levels, and (**F**) hepatic cholesterol levels. (**G**) mRNA expression of cholesterol metabolism genes expressed. as fold change relative to HKDC1^fl/fl^. Total bile acids were quantified in (**H**) serum and (**I**) liver. (**J**) Untargeted metabolomics was used to measure levels of bile acid species. (K) Feces were collected over 24 hours and used to quantify fecal bile acids. N = 3-5 for each genotype per condition. Data were analyzed using one-way ANOVA with Dunnett correction for I, K, two-way ANOVA with Šidák correction for A -G, and *t*-test for H-I. *p < 0.05, **p < 0.01, ***p < 0.001.

Hepatic bile acid metabolism is disrupted in MASLD, and excessive bile acid synthesis and signaling can exacerbate metabolic dysfunction, promoting steatosis and potentially accelerating progression to MASH. In addition, while both serum and hepatic bile acid levels were reduced (Fig 6H–I), metabolomics data revealed a significant increase in hepatic muricholic acid (MCA) levels (Fig 6J), which is the primary hydrophilic bile acid in mice and is known to enhance the fecal excretion of cholesterol. Supporting this observation, an increase in fecal bile acids was also noted, although this was not statistically significant (Fig 6K). The expression of Cyp7A1—the rate-limiting enzyme in the classical bile acid synthesis pathway—was enhanced (S Fig 2C), a change previously associated with inhibiting MASH progression. With decreased systemic and hepatic cholesterol levels, reduced bile acid levels can inhibit MASH progression by decreasing liver inflammation, oxidative stress, and lipid accumulation—key drivers of liver damage in MASLD. Since mice had no difference in food intake or energy expenditure but still gained less adiposity, we reasoned gut lipid absorption might be affected. To assess this, we quantified fecal NEFAs in the WD-fed mice and found that although there was no change in fecal NEFAs at the start of the study (S Fig 2D), there was a significant increase in fecal NEFAs towards the end of the study (S Fig 2E). This observation points to a change in the absorption of lipids in the gut, and since HKDC1 has high expression in the gastrointestinal tract, we assessed the expression of HKDC1 in the ileum and found no changes (S Fig 2F). We also quantified the expression of some inflammatory and metabolic genes in the ileum and found no significant difference (S Fig 2F).

### Transcriptomic analysis reveals that hepatic HKDC1 ablation inhibits pro-inflammatory and pro-fibrogenic pathways

To further analyze how hepatic HKDC1 knockout ameliorates diet-induced MASH progression, liver samples from HKDC1^LKO^ and HKDC1^fl/fl^ mice were subjected to bulk RNA sequencing. Compared to the HKDC1^fl/fl^ livers, 93 genes were upregulated, and 379 genes were downregulated in HKDC1^LKO^ livers (Figure 7A-B). Gene ontology analysis of the downregulated genes revealed that the “extracellular matrix organization” pathway is the most affected pathway (Figure 7C). A further in-depth analysis of this pathway revealed that most of the genes belonged to the pro-fibrogenic program (Fig 7D). Multiple other pathways related to the extracellular matrix or cell adhesion were also enriched, including “cell-substrate adhesion”, “wound healing”, “cell-matrix adhesion” (Figure 7C) and “cytokine-mediated signaling pathways” (Fig 7E). These pathways are consistent with our finding that hepatic HKDC1 knockout reduces hepatic inflammation and fibrosis (Fig 5I-K). We also performed GSEA using the whole gene list and their log_2_ fold change. In addition to the pathway mentioned above, GSEA also showed that pathways related to fatty acid metabolism were enriched in the female HKDC1^LKO^ liver sample, which is in line with reduced steatosis in mice livers with HKDC1 deficiency (Supplementary Figure 3A-B). In summary, bulk RNA-seq confirmed that hepatic HKDC1 ablation protects the mice from WD-induced steatosis, inflammation, and fibrosis. We also performed a widely targeted metabolomics analysis on these samples, but the results showed no significant changes (S Fig 3C). Perhaps a targeted lipidomics analysis will yield more informative results in the future.

**Fig 7:**
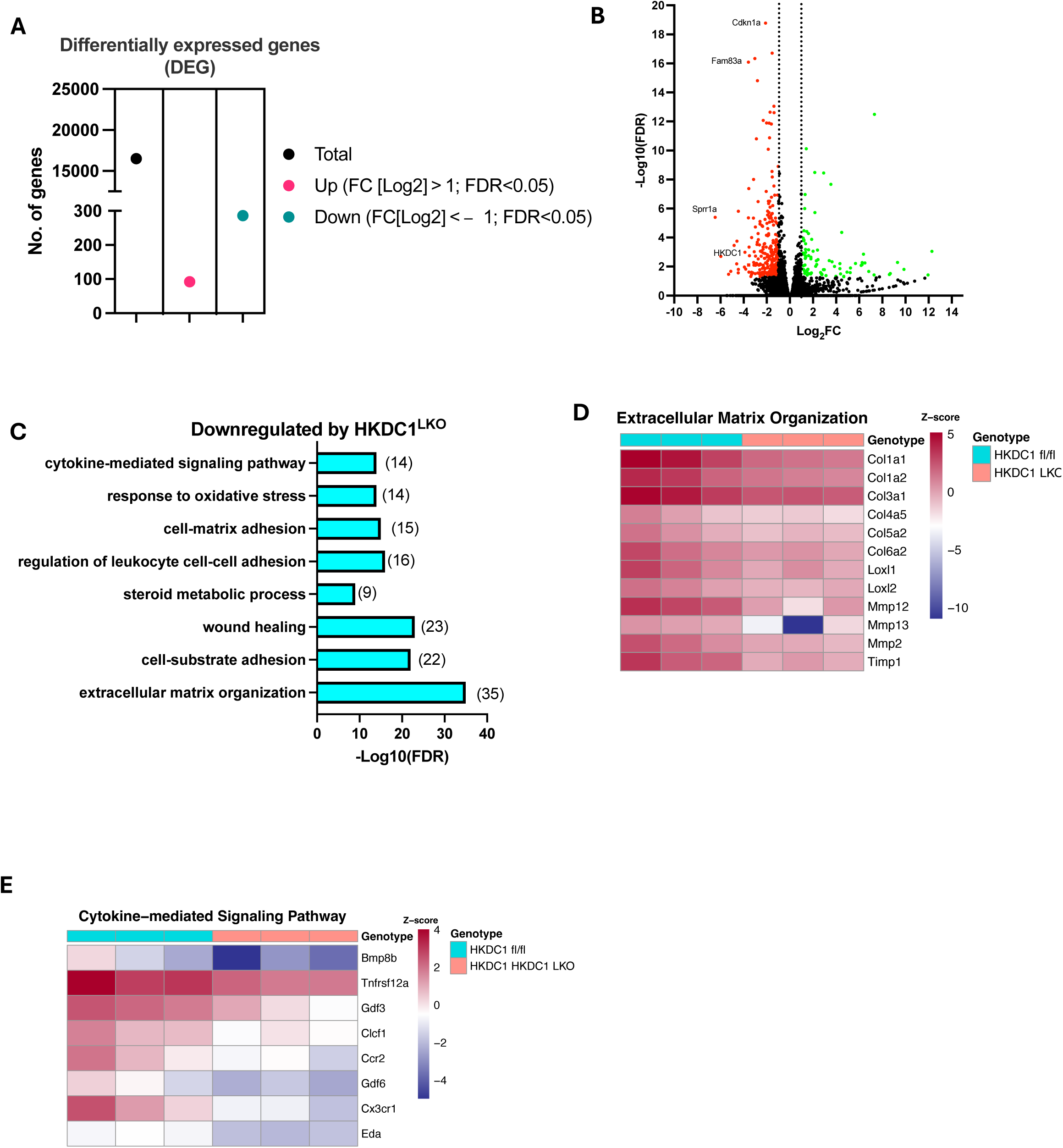
Transcriptome analysis reveals that hepatic HKDC1 deficiency is enriched in pathways that protect against WD-induced MASH. Total mRNA was extracted from liver tissues and subjected to bulk mRNA sequencing. (**A**) Total differentially expressed genes, (**B**) volcano plot of upregulated and downregulated genes, (**C**) Gene ontology analysis was performed to show pathways downregulated in HKDC1^LKO^, heatmaps for (**D**) genes from “extracellular matrix organization” and (**E**) cytokine mediated signaling. N = 3 for each genotype per condition.

### Targeted Ablation of Hepatic HKDC1 Drives Gut Microbial Changes that Inhibit MASH Progression

MASH progression is closely linked with gut microbiome dysbiosis. To evaluate this possibility in our model, we assessed gut microbial composition by 16S rRNA sequencing of the cecal samples from HKDC1^LKO^ and HKDC1^fl/fl^ mice at the terminal time. The relative abundance of each phylum is shown in Figure 8A. No differences were observed in richness (Chao1; Fig 8B) and diversity (Simpson and Shannon; Fig 8C-E) within each sample for [1] diversity. The significantly changed gut microbiota at different levels using Metagenome-seq are shown in Table 2. *Coriobacteria, Erysipelotrichaceae, Faecalibaculum,* and *Bacteroidaceae* previously positively associated with MASH (green font), had higher counts in the HKDC1^fl/fl^ mice^45–48^. *Listeriaceae,* a family vulnerable to bile acid, was higher in the HKDC1^LKO^, possibly due to reduced bile acid secretion into the gut in the HKDC1^LKO^ mice^49^. Bacillus at the genus level (red font) were enriched in female HKDC1^LKO^ mice, which has been shown to alleviate MASH progression when used as supplementation^50,51^. Linear discriminant analysis (LDA) Effect Size identified *Lactobacillales* as a biomarker for less MASH progression in female HKDC1^LKO^ mice (Figure 8F). *Lactobacillus* treatment has also been shown to alleviate MASH-induced pathogenesis^52^. In summary, the gut microbiome of HKDC1 liver-specific knockout mice has fewer bacterial strains that positively correlate with MASH, which may partially contribute to or result from the improved phenotype of mice with hepatic HKDC1 ablation.

**Fig 8:**
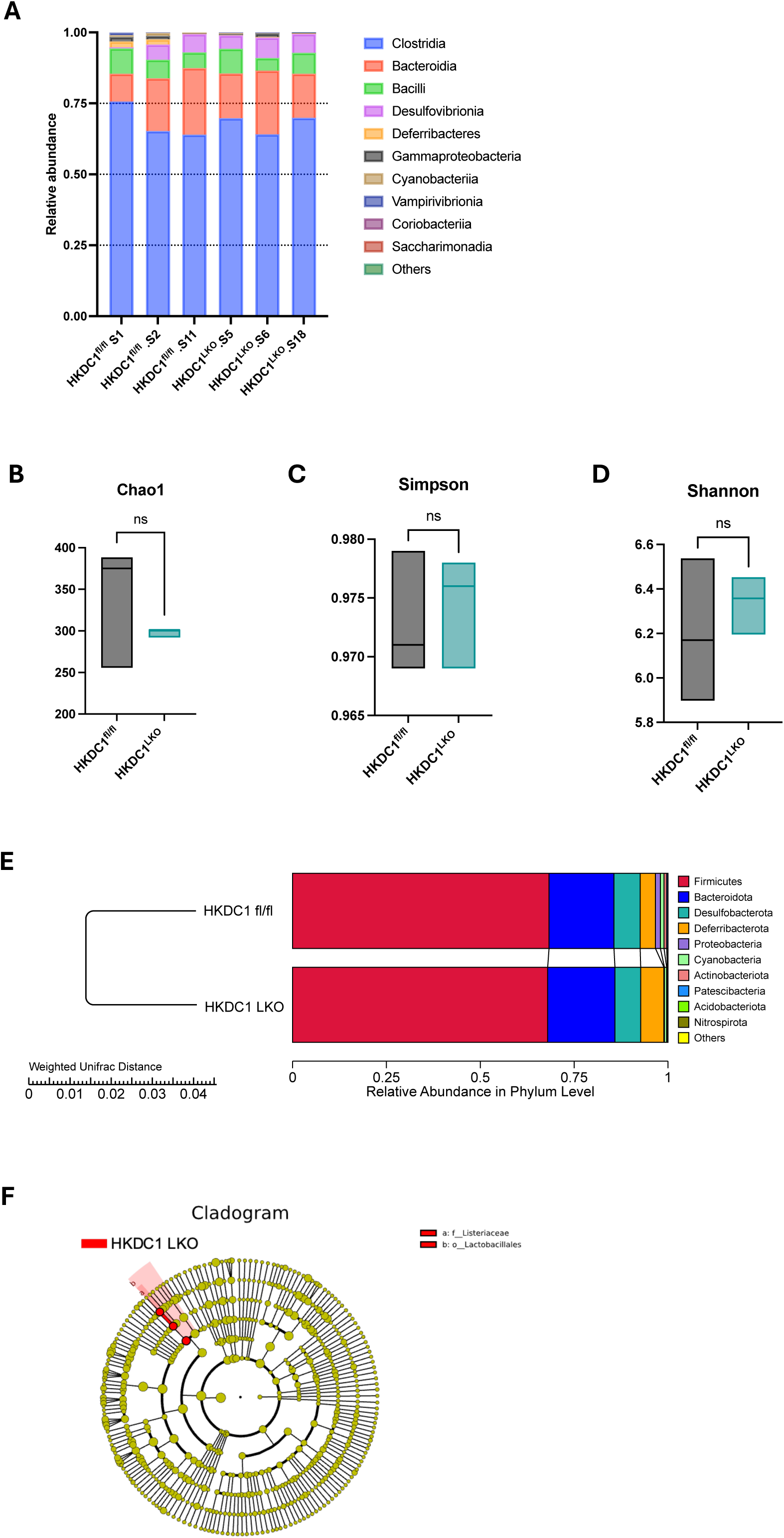
The gut microbiome of hepatic HKDC1 deficiency mice possesses MASH-antagonizing gut bacterial strains. Twenty weeks WD fed female HKDC1^fl/fl^ and HKDC1^LKO^ mice cecal contents were subjected to 16S rRNA sequencing. (**A**) The relative abundance, (**B**) l1 diversity indices including (**C**) Chao1 and (**D**) Shannon index). (**E**) β diversity was analyzed using the Unweighted Pair-group Method with Arithmetic Mean (C; UPGMA), Cladogram of LEfSe (linear discriminant analysis (LDA) Effect Size) analysis. (**F**) significantly different species richness from Metagenomic sequencing. N = 3 for each genotype per condition.

**Table 2:**
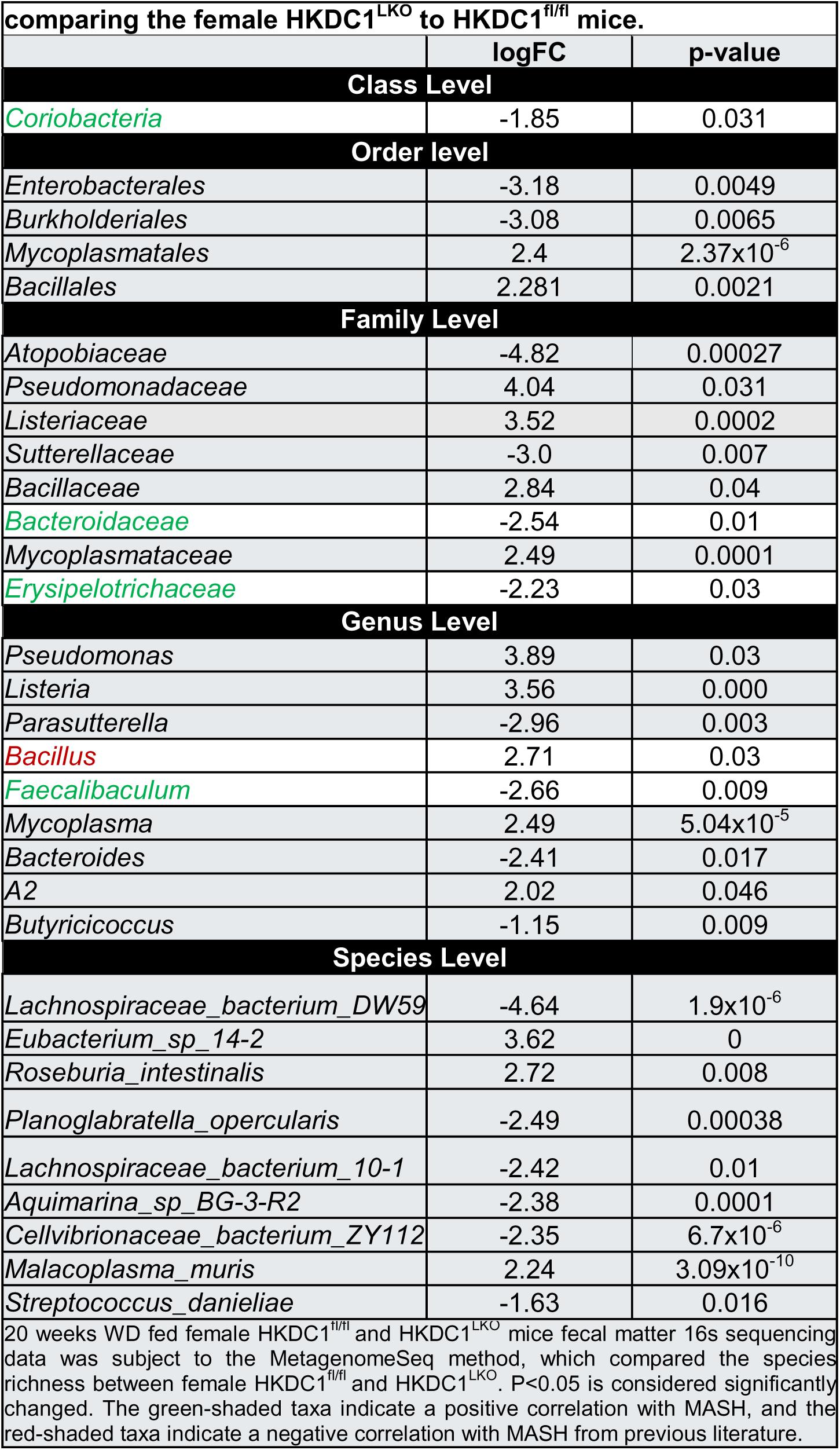
Significantly changed taxa of the fecal microbiota when comparing the female HKDC1LKO to HKDC1fl/fl mice.

## Discussion

MASH has become a health concern as its global prevalence rate is 30%^53^. It will likely worsen as food availability improves, owing to economic growth in developing countries^2,4,5^. Women are more likely to be affected by MASH, especially after menopause^8–12^. Currently, apart from calorie restriction and the just approved resmetirom^54,55^, there are limited therapeutic strategies available for MASH patients. Therefore, identifying novel genes that can potentially be targeted is crucial. HKDC1 is a novel metabolic protein, and our work over the past five years has put it in the limelight in human pathological conditions, particularly in the liver. Our group was the first to show that HKDC1 has negligible expression in adult healthy hepatocytes^56^ and is upregulated in pathophysiology, such as MASLD in male mice^42^ and liver cancer^30^. Our recent work also showed that targeting HKDC1, particularly its mitochondrial interaction, can be beneficial in the fight against liver cancer^57^. Our current work describes HKDC1 as a crucial player in the progression of diet-induced MASH in female mice for the first time. By examining HKDC1 expression and function in the context of metabolic stress, we seek to uncover its contributions to hepatic glucose and lipid metabolism and its potential as a therapeutic target for mitigating MASH progression.

First, data from our human cohort established a strong positive correlation between upregulated hepatic HKDC1 expression and MASLD progression in females. Second, data from our mouse cohort highlights the molecular effects of hepatic HKDC1 as a potential regulator of lipid (cholesterol-bile acid) metabolism in MASLD progression in females. Third, knockout of hepatic HKDC1 in NC-fed female mice did not alter glucose tolerance, hepatic TG, cholesterol, and serum ALT. This evidence indicates that hepatic HKDC1 may be dispensable in a healthy state. Earlier research shows that HKDC1 is scarcely expressed in a healthy state^26^. Lastly, from our mouse cohort data, hepatic HKDC1-dependent gut-liver axis modulating intestinal fatty acid absorption also emerges.

After the breakthrough Hyperglycemia and Adverse Pregnancy Outcomes (HAPO) Study associating *HKDC1* genetic variants with pregnancy glycemic traits^44^, several meta-analyses and GWAS studies have linked *HKDC1* with women’s health, especially during pregnancy^58^ and polycystic ovary syndrome^59^. However, questions remained regarding the functional role of HKDC1 in women’s health. Through the combined use of clinical and preclinical models, here we have provided insights into the role of HKDC1 in the human female liver: a direct relationship with the nutrient (over)load. Though not a stand-alone observation and reiterated earlier in male human cohorts^16,26^ and extrahepatic tissues like enterocytes in mice^60^, it re-emphasizes the unique sensitivity of HKDC1 to nutrients compared to other HKs. Factors governing this temporal expression of HKDC1 during the course of the disease remain elusive. Sex hormones may play a role based on their increased hepatic expression during pregnancy in mouse livers^28^ and up-regulation of androgen receptor regulon in female MASH mice^61^. While this indicates a possibility of HKDC1’s role in sex-dependent disparity in MASLD incidence, conclusive data can be obtained only after comparing and contrasting male and female HKDC1^LKO^ MASH and gonadectomized mouse models. Notably, our female mice (∼36 weeks of age at the end of the experimental period) would not have reflected menopausal phenotype, again underscoring the nutrient excess-HKDC1 expression link.

As follows, HKDC1 ablation in the liver restricted WD-induced increase in the body weight and fat mass and reduced fat accumulation in the liver. This phenotype is protective as MASH livers exhibit dysregulated glucose production, lipid storage, and metabolism^45,62^. Lipid burden in MASH livers can arise from disruption in lipid supply (increased absorption of circulating lipids) and lipid elimination (lipid oxidation/ secretion). Our bulk RNA seq data revealed that fatty acid metabolism pathways were enriched in HKDC1^LKO^ mice livers. mRNA expression of several genes of fatty acid oxidation, *Ppara, Cyp4A14, Cpt1a, and Cpt2,* showed only a modest, insignificant increase in HKDC1^LKO^ livers with a significant increase in *Ppara* responsive gene *Cyp4A10*. Fatty acid oxidation (microsomal ω-hydroxylation) enzyme Cyp4A10 has been associated with diet-induced steatosis and inflammation^63,64^, though it is also known to be classically induced in response to fasting^65^. Since this increase in Cyp4A10 expression was without inflammation and fibrosis in HKDC1^LKO^ livers, it suggests an increased hepatic lipid turnover contributing against WD-induced MASH development. Increased Fabp1 mRNA expression further indicates altered intrahepatic trafficking of fatty acids.

HKDC1 knockout reduces systemic and hepatic cholesterol, a finding that may result from decreased synthesis, increased efflux, or a combination of both. In support of this, our observations of altered expression levels of key cholesterol metabolism genes such as *Hmgcr, Ldlr,* and *Cyp7a1* suggest integrated changes in hepatic lipid regulation in HKDC1^LKO^ livers. To pinpoint the underlying mechanisms, in future studies, it will be crucial to 1) quantify the rate of hepatic cholesterol synthesis, 2) assess whether and how HMGCR—the rate-limiting enzyme in cholesterol synthesis—is being downregulated, and 3) determine if enhanced cholesterol efflux is preventing its accumulation in hepatocytes. Studies have also shown that elevated MCA levels promote increased fecal excretion of cholesterol and free fatty acids by decreasing lipid absorption via multiple mechanisms^66–68^: glycine-β-MCA treatment enhances fecal bile acid excretion and reduces gut bile acid hydrophobicity, higher MCA levels reduce cholesterol uptake from the portal vein, and increased MCA concentrations boost hepatic cholesterol exporter expression^66^. Collectively, these data suggest that HKDC1 ablation reduces hepatic cholesterol accumulation by decreasing cholesterol synthesis and promoting MCA production to enhance cholesterol excretion, thereby inhibiting MASH pathogenesis (Fig 9). However, since hepatic bile acid synthesis only contributes to about 5% of the total bile acid pool, it would be premature to conclude that HKDC1 ablation reduces the overall bile acid pool. A comprehensive analysis—including measurements from blood, liver, gall bladder, intestine, and feces—is needed to establish a detailed baseline understanding of how HKDC1 ablation affects total bile acid pools and composition. In the future, it will be important to thoroughly examine the enterohepatic signaling that governs fatty acid and bile acid metabolism in this model, especially given the markedly reduced intestinal absorption of fatty acids observed in HKDC1LKO mice. The reduced gut fatty acid absorption may be secondary to hepatic HKDC1 ablation due to bile acids’ role in fat emulsion and absorption^69^, as well as cholesterol’s role in bile acid synthesis; however, this possibility requires further exploration. Additionally, combining pharmacologic and/or genetic modifications of fatty acid metabolism genes with lipidomics and single-cell RNA sequencing will be crucial to fully understand the interaction of HKDC1 with these components in regulating hepatic lipid metabolism in MASH. With this study, we have taken initial steps toward elucidating the pathways downstream of hepatic HKDC1 ablation that may contribute to the previously noted reductions in triglycerides and cholesterol in our mouse models^28,29^, positioning hepatic HKDC1 as a promising target for MASH treatment, particularly in women.

**Fig 9.**
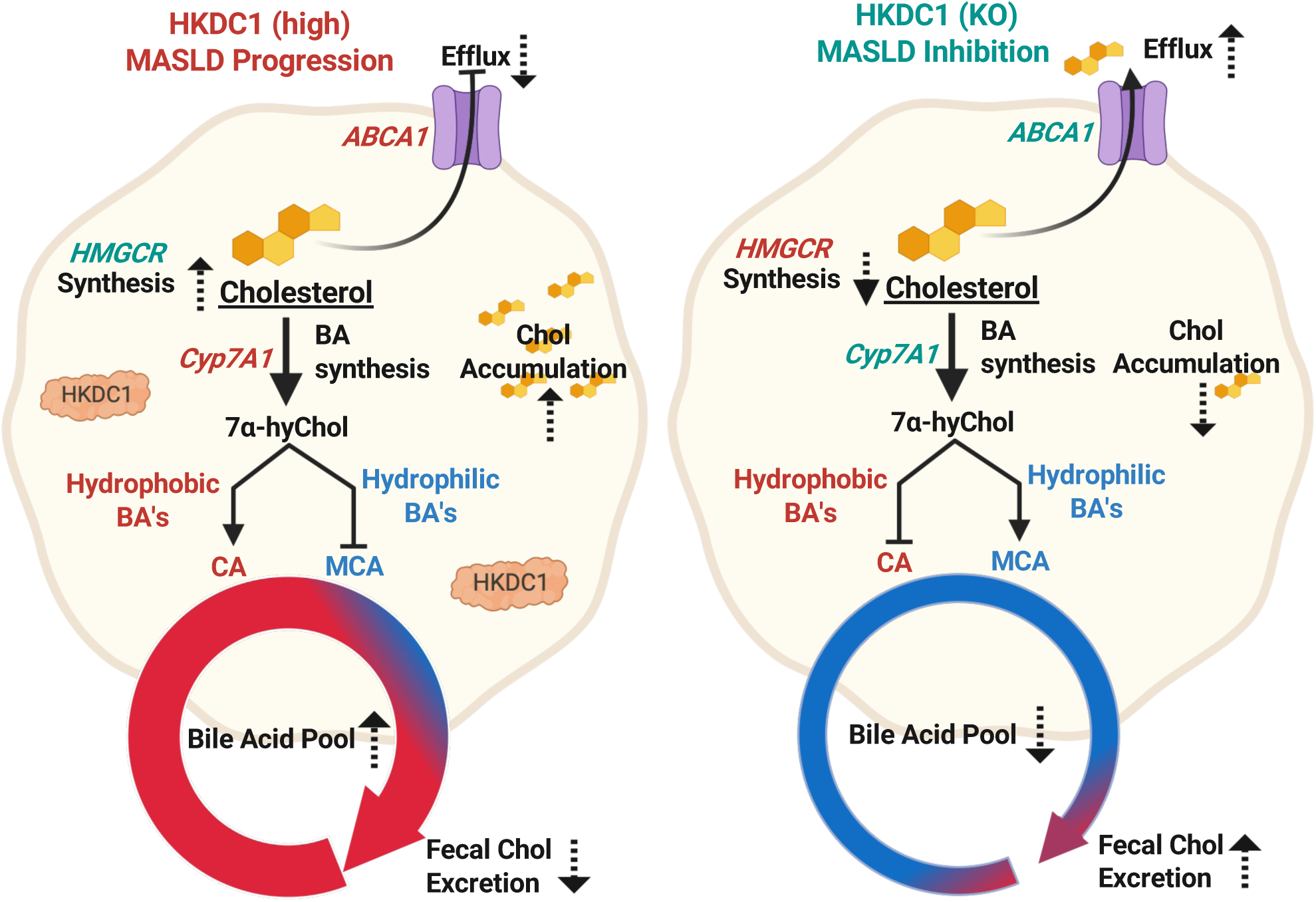
HKDC1 in MASLD – The song of Chol & Bile. This schema illustrates the potential roles of HKDC1 in cholesterol and bile acid metabolism during the progression of MASH. When HKDC1 is high (left), cholesterol synthesis is upregulated, and bile acid pools consist mainly of hydrophobic species (CA; red), which results in cholesterol accumulation within the hepatocytes. These factors collectively contribute to the progressive pathology of MASH. When HKDC1 is ablated (right), HMGCR expression decreases, thereby decreasing endogenous cholesterol synthesis. Meanwhile, there is a shift in bile acid synthesis favoring the production of muricholic acid (MCA; blue). MCA enhances both the efflux of cholesterol via ABCA1 and excretion in feces. Upregulated genes are in green, and downregulated genes are in red. The dotted arrows indicate the increase or decrease in processes where we hypothesize HKDC1 exerts its influence. Created in Biorender.

These findings have broader implications beyond MASH, particularly in other metabolic diseases where sex differences are critical. Given that HKDC1 has been linked to pregnancy-related glucose regulation, polycystic ovary syndrome (PCOS), and insulin resistance, its role in broader metabolic dysfunction warrants further investigation. Metabolic diseases such as type 2 diabetes, obesity, and cardiovascular disease (CVD) exhibit sex-specific differences in prevalence, progression, and response to treatment, which could be influenced by differential HKDC1 expression. For example, the observed connection between HKDC1 and lipid metabolism may extend to atherosclerosis and dyslipidemia, conditions that disproportionately affect postmenopausal women due to hormonal changes. Additionally, the gut-liver axis regulation by HKDC1 suggests potential interactions with the gut microbiome and enterohepatic signaling, which could be relevant for disorders such as metabolic syndrome and inflammatory bowel diseases. Future studies should explore whether sex hormones directly regulate HKDC1 expression and how its inhibition might affect systemic metabolism across different metabolic conditions. Understanding these mechanisms could pave the way for sex-specific therapeutic strategies targeting HKDC1, optimizing treatment approaches for both MASH and other metabolic disorders.

### Future Directions and Therapeutic Interventions Targeting HKDC1

Future studies should focus on therapeutic interventions targeting HKDC1, particularly for MASH and related metabolic disorders, to translate these findings into clinical applications. One promising approach is the development of small-molecule inhibitors to selectively reduce hepatic HKDC1 activity, thereby lowering lipid burden and improving metabolic homeostasis. Given HKDC1’s mitochondrial interactions, targeting HKDC1-dependent metabolic pathways, such as fatty acid oxidation and cholesterol metabolism, could provide therapeutic benefits in preclinical models of MASH. Additionally, gene therapy strategies using CRISPR-based gene editing or siRNA approaches could be employed to modulate HKDC1 expression in a tissue-specific manner, with AAV-mediated HKDC1 silencing offering insights into long-term metabolic effects. Another emerging strategy is using PROTACs (Proteolysis Targeting Chimeras), which could enable targeted degradation of HKDC1, providing a more precise therapeutic approach than traditional inhibitors. Furthermore, combination therapies should be explored, such as HKDC1 inhibition alongside FXR agonists to enhance cholesterol-bile acid homeostasis or in conjunction with resmetirom (THR-β agonist) to assess its impact on lipid metabolism and fibrosis reduction. Finally, analyzing human liver biopsies from MASLD/MASH patients will be crucial to determining whether HKDC1 expression correlates with disease severity, fibrosis progression, and treatment outcomes, ultimately paving the way for precision medicine approaches targeting HKDC1 in metabolic diseases.

