## Supplementary figures and images for "Hepatic HKDC1 Deletion Alleviates Western Diet-Induced MASH in Mice"

### Supplemenatry Figures

##### Supplementary Figure 1

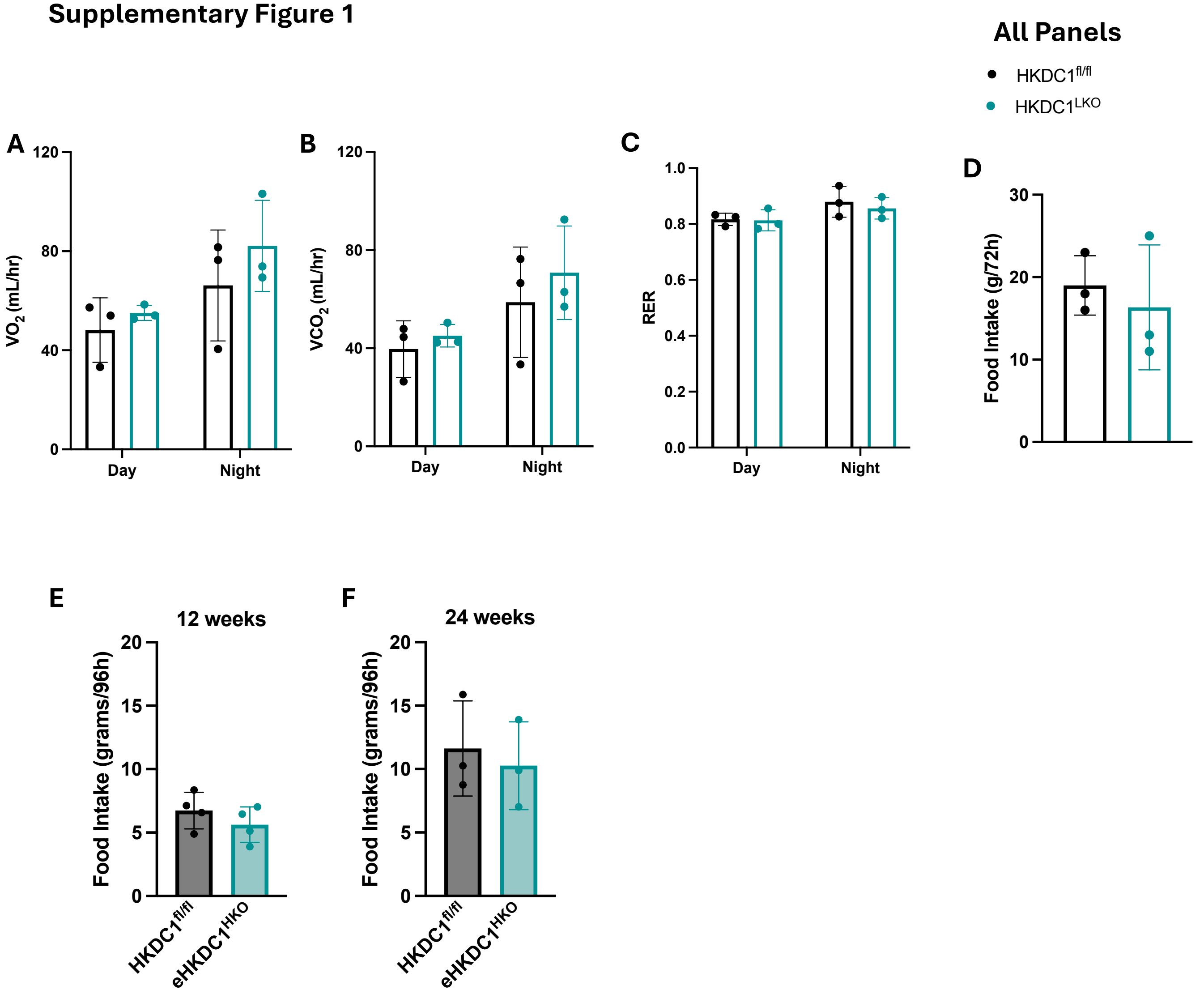

##### Supplementary Figure 2

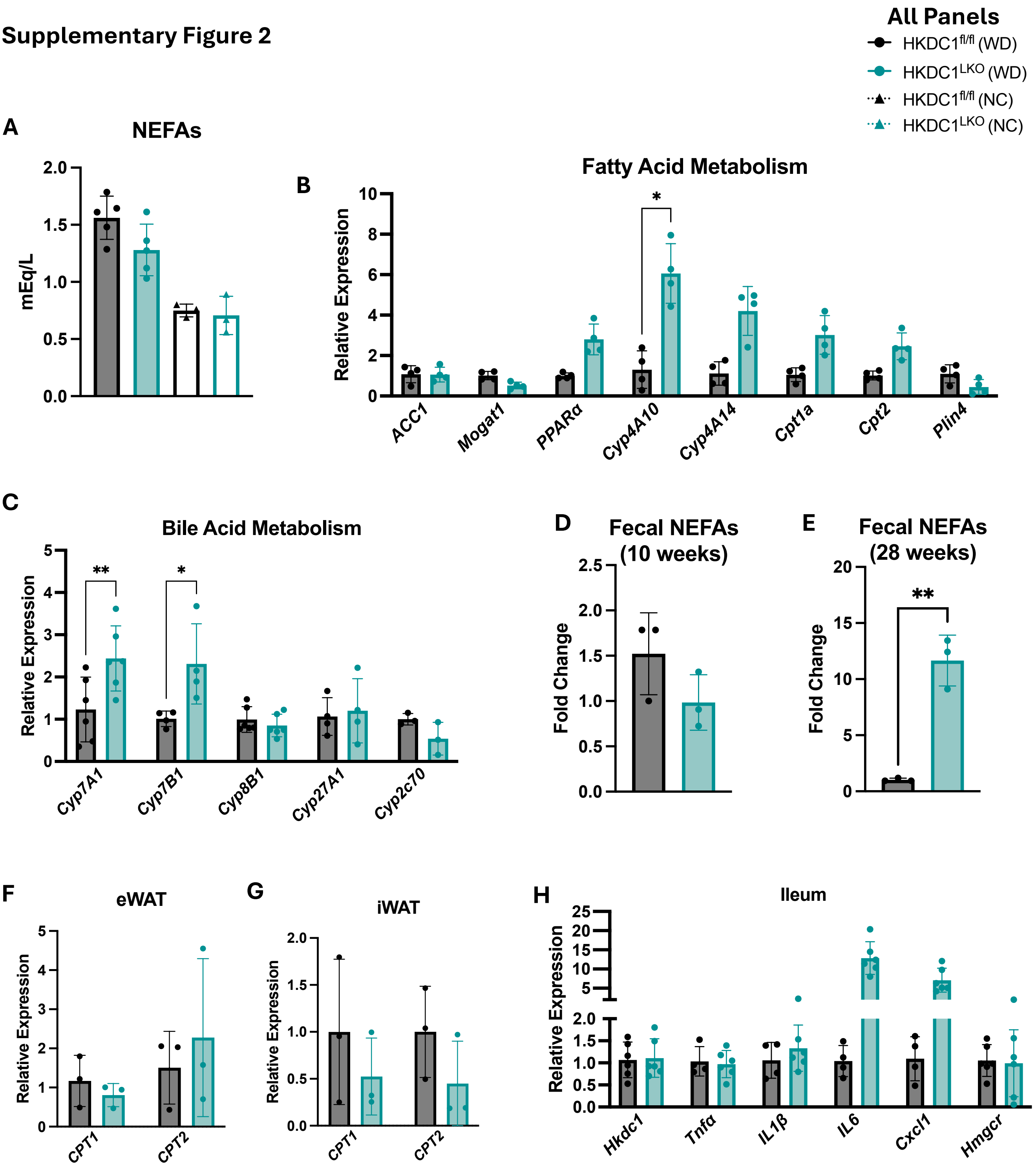

#### Supplementary Figure 3

A

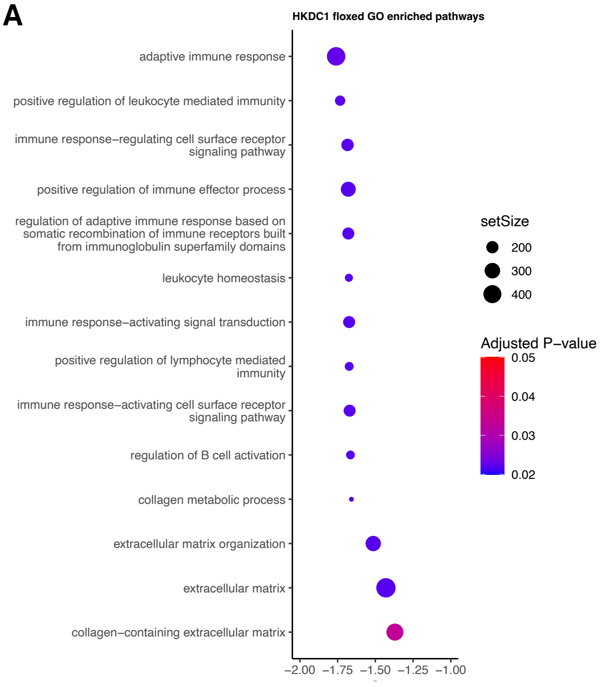


B

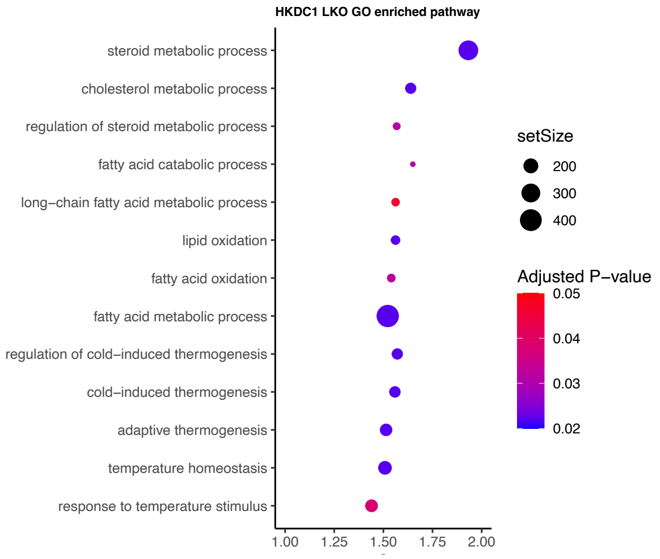


C

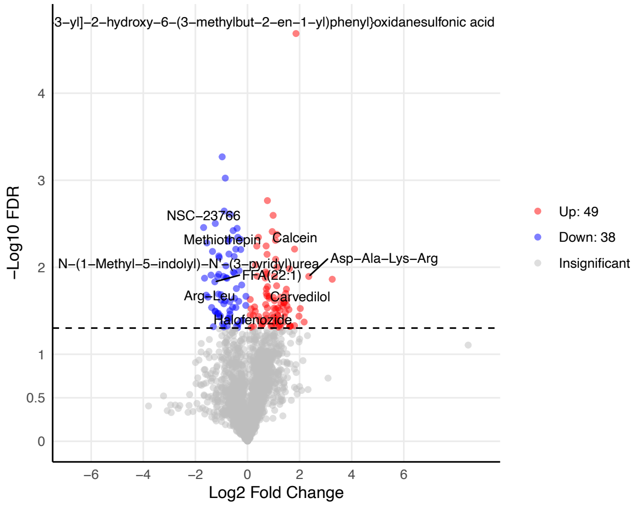
